# Genomic insights into the adaptive evolution of the memorial rose (*Rosa lucieae*; Rosaceae) in saline environments

**DOI:** 10.1101/2024.08.29.610403

**Authors:** Ji-Hyeon Jeon, Myunghee Jung, Younhee Shin, Seung-Chul Kim

**Affiliations:** Department of Biological Sciences, Sungkyunkwan University, Suwon 16419, Republic of Korea; Research and Development Center, Insilicogen Inc., Yongin 16954, Republic of Korea

**Keywords:** *Rosa lucieae*, *Rosa wichurana*, rose genome, salt tolerance, salinity resistance, adaptive evolution

## Abstract

*Rosa lucieae* (syn. *Rosa wichurana* or *R. wichuraiana*; Rosaceae) is one of the major wild progenitors of modern rose cultivars. In breeding roses, *R. lucieae* has contributed not only to white blooms and shiny leaves, but most of all, to succeed in various harsh conditions and high resistance to diseases and pests. The wild *R. lucieae* inhabits littoral sand banks or rocks, where drought stress and high salinity inhibit plant growth. In this study, we assembled the chromosome-level genome sequence of a wild accession of *R. lucieae* and identified potential genes that may responsible for the evolution of salt tolerance. Within the 544.7 Mb-long genome of *R. lucieae*, recent duplications and divergence of the genes responsible to stress tolerance were identified, including stress response, signaling, homeostasis, and detoxification. Additionally, the gene family evolution analyses elucidated rapid evolution of stress tolerant functions in *R. lucieae*. The first high-quality genome assembly of wild *R. lucieae* contributes significantly to our understanding of the genomic evolution and adaptive mechanism of *R. lucieae*, which promises the enhancement of our efforts in rose breeding programs.

## INTRODUCTION

The roses are the flowering plants belonging to the genus *Rosa* in the family Rosaceae, encompassing 100–200 wild species and more than 30,000 cultivars (Rehder, 1940; Young et al., 2007). As renowned for their beauty and fragrance, roses have been beloved for millennia along with the history of human civilization, and cultivated for ornamental, aromatizing, therapeutic and culinary purposes in global (Rehder, 1940; Yu and Ku, 1985; Gudin, 2003; Nybom, 2009). Despite importance of roses, rose cultivation has faced various hurdles to overcome, including environmental changes, diseases and pests, hybrid incompatibility, and productivity (Gerard, 1897; Van Fleet, 1908; Gudin, 2003; Nybom, 2009). However, breeding hardy wild roses developed many robust hybrid cultivars which can cope with those problems (Van Fleet, 1908; Gudin, 2003). *Rosa lucieae*, also well known as its synonym, *Rosa wichurana* or *R. wichuraiana,* or the memorial rose, has been cherished for centuries as one of the major progenitors contributed to developing many modern rose cultivars (Gerard, 1897; Van Fleet, 1908; Yu and Ku, 1985; Crespel and Mouchotte, 2003; Marriott, 2003; Nybom, 2009). *Rosa lucieae* is a rambler rose vigorously growing even in the bad climate and on bare soil or rocks (Gerard, 1897; Rehder, 1940; Yu and Ku, 1985), highly resistant to the prevalent rose diseases such as black spot (*Diplocarpon rosae*; Noack, 2003; Byrne et al., 2005), and easy to cross (Gerard, 1897; Van Fleet, 1908). Such characteristics of *R. lucieae* have led the successful development of thousands of robust *R. lucieae* hybrid cultivars (Gerard, 1897; Van Fleet, 1908; Rehder, 1940; Yu and Ku, 1985; Byrne et al., 2005). Consequently, in the classification scheme of modern roses, the Hybrid Wichurana is now identified as one of the cultivar groups, bred by crossing *R. lucieae* and other roses and characterized by its rambling habit and wide variety of flower shapes and colors (Cairns, 2003).

*Rosa lucieae* has been classified as a member of the section *Synstylae* within genus *Rosa*, where the species are closely related but diversified with unique characteristics to adapt to various ecological habitats (Rehder, 1940; Wissemann, 2003; Jeon et al., 2024). Unlike most of other *Rosa* sect. *Synstylae* species, the natural habitats of *R. lucieae* are open and exposed rocky slopes or banks, especially along the sea shores and riverside at low elevation (Rehder, 1940; Ohba, 1993; Ku and Robertson, 2003). In these habitats, both abiotic stresses such as high light, high salinity and heavy metals, and biotic stresses caused by various herbivores and pathogens restrict plant growths and introduction. Nevertheless, *R. lucieae* has adapted to harsh environments of those habitats, evolving to resist and tolerate to various abiotic and biotic stresses. In addition to their ornamental characteristics, stress resistance and resilience in various environmental conditions are pivotal in breeding roses, efficiently increasing productivity and durability (Van Fleet, 1908; Gudin, 2003; Debener and Linde, 2009; Leus et al., 2018). Utilizing wild roses including *R. lucieae* in the rose breeding programs will pave the way for introducing the stress tolerant traits to cultivated roses (Van Fleet, 1908; Noack, 2003; De Cock et al., 2007). Understanding genetic and genomic architectures of roses also promises improvement in breeding roses for the resistance by allowing the applications such as marker-assisted selection and genetic engineering (Debener and Hibrand-Saint Oyant, 2009; Debener and Linde, 2009; Gahlaut et al., 2021). Recently, several transcriptomic approaches illustrated genetic functions and mechanisms of the stress responses in hardy wild roses, including disease resistance (Xiang et al., 2019) and cold tolerance (Zhang et al., 2016; Zhuang et al., 2021). Additionally, the genome of *Rosa rugosa* provided insights into the evolution of stress response in harsh environments (Chen et al., 2021). Nevertheless, the genomic intricacies of roses have impeded the extensive understanding on the genomic bases of key characteristics in the rose cultivation (Gahlaut et al., 2021).

The salt stress is one of the major abiotic stresses in plants, which induces osmotic stress, ion toxicity, oxidative stress, and decreasing in photosynthesis (Zhu, 2001; Munns and Tester, 2008). In agriculture, nearly 20 % of global irrigated lands have been salinized, and recurrent irrigation with brackish groundwater or return flow caused from limited sources of fresh water accumulates salts in agricultural lands (Ghassemi et al., 1995; Tilman et al., 2002; Munns, 2005). Meanwhile, the increased level of soil salinity has severely affected the decline in productivity and quality of crops, resulting in consistent and complex economic losses (Munns and Gilliham, 2015). Not only can the high concentration of salts directly affect to the cytological and physiological dysfunction, but also involve other secondary cytotoxic effects caused from generation of reactive oxygen species (ROS), imbalance of other essential ions such as K^+^, and malnutrition (Zhu, 2001; Botella et al., 2005). In plants, three major mechanisms of salt tolerance are known: homeostasis, detoxification, and growth control (Zhu, 2001; Munns, 2005; Munns and Tester, 2008). Homeostasis can be established by compartmentalization of Na^+^ and Cl^−^ into the vacuoles, modulation Na^+^ influx and efflux, and accumulating compatible osmolytes, which are regulated by root endoderm and xylem, or stress signaling pathway including Ca^2+^-mediated cascades (Zhu, 2001; Botella et al., 2005; Munns, 2005; Van Zelm et al., 2020). Ionic and oxidative toxicity can be detoxified or prevented by protein protection and repair, antioxidant metabolites, and mitogen-activated protein kinase (MAPK) cascades-induced detoxifying enzymes (Zhu, 2001; Chinnusamy and Zhu. 2003; Botella et al., 2005; Munns, 2005; Van Zelm et al., 2020). Salt stress also can inhibit the plant growth with stomal closure induced by abscisic acid (ABA), restricting the photosynthesis (Botella et al., 2005; Van Zelm et al., 2020). However, the growth-promoting hormones such as jasmonic acid (JA), brassinosteroid (BR), and gibberellic acid (GA) are responsible for the growth recovery after salt stress (Van Zelm et al., 2020).

Roses are generally salt sensitive with lower yield, fewer flowers, and poor growths even in relatively low salt concentrations (Cabrera and Perdomo, 2003; Cabrera et al., 2009; Niu and Sun, 2017). Nevertheless, the genetic or genomic studies of roses not only lag behind other rosaceous crops (e.g., apples, peaches, and strawberries; Shulaev et al., 2008; Foucher, 2009; Gahlaut et al., 2021), but the salt tolerance of roses is also poorly understood compared to the aesthetic features (e.g., flower forms, colors, and fragrance), which have been priorly considered in breeding roses (Gudin, 2003; Bao et al., 2021). Given the deteriorating soil salinity stemming from recurrent irrigation, urbanization and climate changes, the salt tolerance becomes pivotal in the rose cultivation (Cai et al., 2014; Corwin, 2021).

Consequently, the advent of high-throughput next-generation sequencing technique has led recent multi-omics investigation into the salt tolerance in various rose cultivars of *Rosa chinensis* (Tian et al., 2018; Bao et al., 2021), *R. rugosa* (Chen et al., 2021; Qi et al., 2023), *R.* × *damascena* and *R.* × *hybrida* (Ren et al., 2024).

In this study, a chromosome-level genome of a wild accession of *R. lucieae*, inhabiting the saline sea shore, was assembled in order to understand the genomic evolution and nature of salt tolerance in *R. lucieae*. The chromosome-level genome of *R. lucieae* and its evolution in salt tolerance will contribute to our understanding of genomic features and adaptive mechanisms of wild roses, and promise to enhance our efforts in rose breeding programs.

## RESULTS

### Molecular sequencing, genome assembly, and annotation of Rosa lucieae

A total of 374.6 million paired-end short-sequence reads with 56.6 Gbp (123.4× coverage depths of the estimated haploid genome size) were generated through Illumina HiSeq platform, and 2.7 million long-sequence reads with 28.7 Gbp (62.6× coverage depths of the estimated genome size) were generated through PacBio Sequel platform (Supplementary Table S1). Each of two transcriptome samples from the young leaves and flower buds was sequenced into 9.0-Gbp data with 89 million paired-end short-sequence reads through Illumina NovaSeq platform (Supplementary Table S1). The *k*-mer analysis using genomic short reads estimated the haploid genome size of *R. lucieae* at 458.3 Mbp, of which repeat content was 59.61 % (273.2 Mbp) and the heterozygosity rate was 1.35% (Table 1; Supplementary Fig. S1).

**Table 1.**
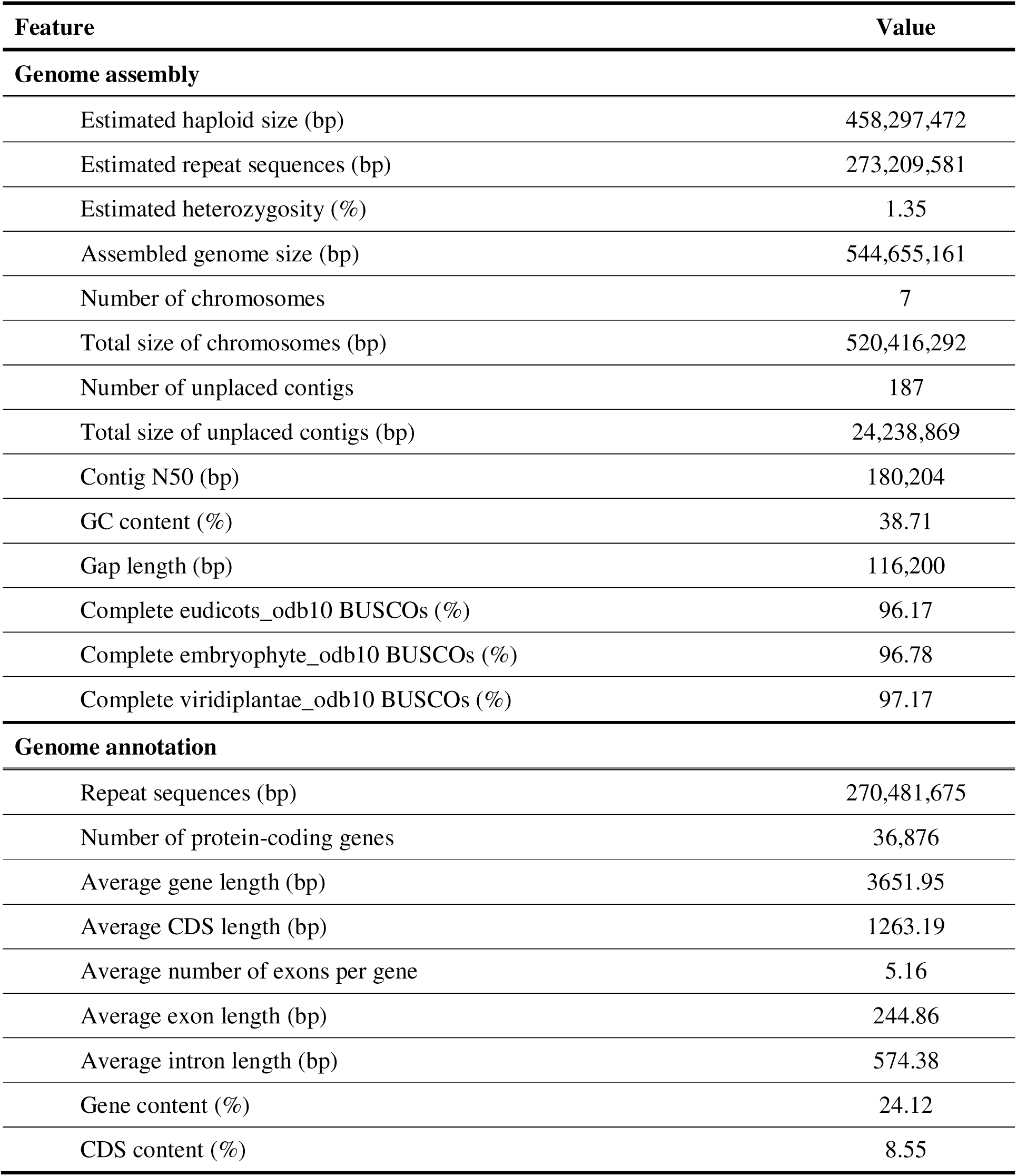
Statistics of the genome assembly and annotation of *Rosa lucieae*.

| Feature | Value |
| --- | --- |
| **Genome assembly** |  |
| Estimated haploid size (bp) | 458,297,472 |
| Estimated repeat sequences (bp) | 273,209,581 |
| Estimated heterozygosity (%) | 1.35 |
| Assembled genome size (bp) | 544,655,161 |
| Number of chromosomes | 7 |
| Total size of chromosomes (bp) | 520,416,292 |
| Number of unplaced contigs | 187 |
| Total size of unplaced contigs (bp) | 24,238,869 |
| Contig N50 (bp) | 180,204 |
| GC content (%) | 38.71 |
| Gap length (bp) | 116,200 |
| Complete eudicots_odb10 BUSCOs (%) | 96.17 |
| Complete embryophyte_odb10 BUSCOs (%) | 96.78 |
| Complete viridiplantae_odb10 BUSCOs (%) | 97.17 |
| **Genome annotation** |  |
| Repeat sequences (bp) | 270,481,675 |
| Number of protein-coding genes | 36,876 |
| Average gene length (bp) | 3651.95 |
| Average CDS length (bp) | 1263.19 |
| Average number of exons per gene | 5.16 |
| Average exon length (bp) | 244.86 |
| Average intron length (bp) | 574.38 |
| Gene content (%) | 24.12 |
| CDS content (%) | 8.55 |

The error-corrected PacBio subreads were *de novo* assembled into 2067 primary contigs with a total of 619.5 Mbp lengths, and this heterozygous primary assembly was curated into the 1356 contigs with a total of 544.9 Mbp lengths. Finally, error sequences of the curated assembly were corrected with short sequence reads, establishing the 544.5 Mbp-long draft *de novo* genome assembly with 1356 contigs (Supplementary Table S2). The BUSCO assessment identified more than 96 % of BUSCO eudicots-, embryophyte-, and viridiplantae-ortholog datasets within the draft assembly of *R. lucieae* respectively (Table 1). The *de novo* assembly was subsequently scaffolded into seven chromosomal sequences and 187 unplaced contigs, finalizing the 544.7 Mb-long chromosomal genomic sequences of *R. lucieae* (Fig. 1; Table 1; Supplementary Table S3). The genome collinearity analysis with the reference chromosomes indicated no genomic gaps, missing data, or possible mis-assembly within the draft genome assembly of *R. lucieae* in the present study (Supplementary Fig. S2).

**Fig. 1.**
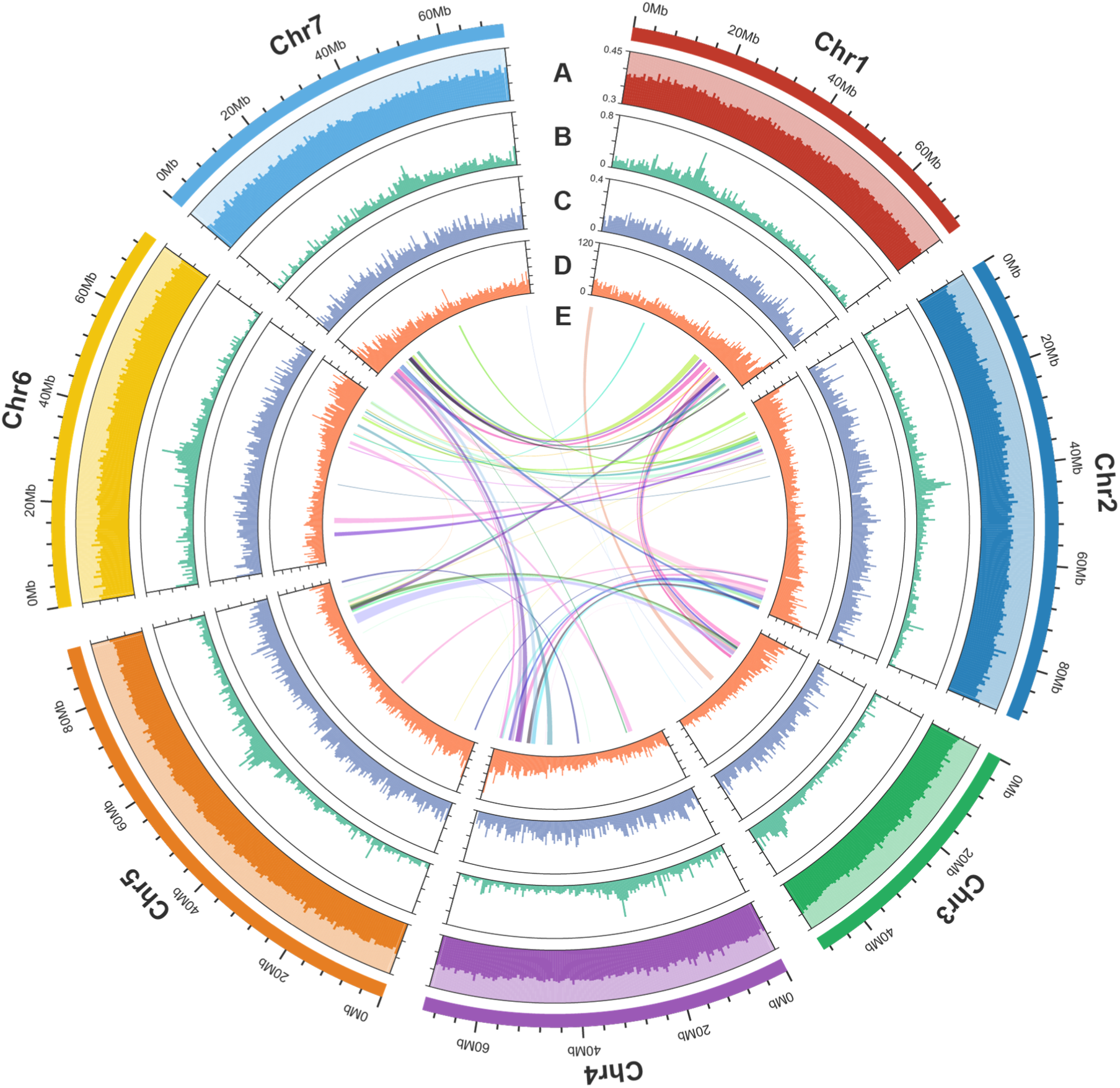
Genome map of *Rosa lucieae*. **A)** GC content, **B)** density of *Gypsy*, **C)** density of *Copia*, **D)** density of protein-coding genes, and **E)** collinear relationships within the genome. The GC content and densities of genetic elements were calculated using sliding windows of 500kb without overlaps.

Within the genome of *R. lucieae*, 49.67 % of sequences (270.5 Mbp) were identified as repeat sequences. Among them, most repeat sequences were interspersed repeats (97.4 %), of which 54.1 % were identified as long terminal repeat (LTR) elements, 15.5 % were identified as DNA elements, and 24.5 % were unclassified (Supplementary Table S4). A total of 36,876 protein-coding genes were identified and annotated within the genome of *R. lucieae* (Table 1), the number of which was close to the numbers of annotated protein-coding gene of other *Rosa* genomes (*R. lucieae*: 32,674; *R. chinensis*: 30,924–39,669; *R. rugosa*: 29,146–39,704 genes; Hibrand Saint-Oyant et al., 2018; Raymond et al., 2018; Chen et al., 2021; Zang et al., 2021; Zhang et al., 2024). Among the protein-coding genes, all the genes except three (36,873 genes; 99.99 %) were functionally annotated from the public databases, including nr, gene ontology (GO), KEGG, and Pfam databases (Supplementary Table S5).

### Structural genomic evolution of Rosa lucieae

The genome of *R. lucieae* was assembled into a total of 544.7 Mbp-long scaffolds and contigs, of which the size was almost similar to the genomes of *R. chinensis* (488.6–541.3 Mbp; sect. *Chinenses*; Raymond et al., 2018; Hibrand Saint-Oyant et al., 2018; Zhang et al., 2024), and slightly to moderately larger than those of *R. rugosa* (401.2–450.0 Mbp; sect. *Rosa*; GCF_958449725.1; Chen et al., 2021; Zang et al., 2021), *R. roxburghii* (504.0–529.9 Mbp; *sect. Microphyllae*; Yang et al., 2024; Zong et al., 2024), and *R. persica* (366.1 Mbp; subg. *Hulthemia*; GCA_025331145.1). The genome size profile of these *Rosa* species was corroborated with the nucleic DNA amount investigation, estimating 2C at 1.13 pg in *R. lucieae*, 1.16 pg in *R. chinensis*, 0.98 pg in *R. rugosa*, 0.95 pg in *R. roxburghii*, and 0.84 pg in *R. persica* (Yokoya et al., 2000). Despite the increased genome size, the repeat content of genome of *R. lucieae* (49.7 %; Table 1; Supplementary Table S4) was nearly similar to *R. rugosa* (50.3–54.1 %; Chen et al., 2021; Zang et al., 2021), whereas it was notably increased in the genomes of *R. chinensis* (61.8–67.9 %; Raymond et al., 2018; Hibrand Saint-Oyant et al., 2018; Zhang et al., 2024).

Within 36,876 protein-coding genes identified within the genome of *R. lucieae*, the ortholog group inference analysis classified 32,666 genes into 19,667 gene family groups, of which 14,436 groups were shared by Rosoideae species (i.e., *R. lucieae*, *R. chinensis*, *R. rugosa* and *Fragaria vesca*), 15,790 groups were shared within the genus *Rosa*, and 586 groups were unique to *R. lucieae* (Fig. 2A). Among 32,666 classified genes, 31,264 were interspecific orthologous and the other 1402 were species-specific paralogous genes (Fig. 2B). Based on the gene annotations of Rosoideae genomes and their homologies, the interspecific collinear blocks were identified to infer the genome synteny. Within the genus *Rosa*, the genome synteny was highly conserved among species, especially between *R. chinensis* and *R. rugosa*. The genome synteny was also moderately conserved between the genera *Rosa* and *Fragaria*, even though several structural variations were identified (Supplementary Fig. S3). Despite the increased repeat content in the genomes of *R. chinensis*, the genome synteny was highly conserved between *R. chinensis* and *R. rugosa*. Moreover, despite the closer phylogenetic relationship of *R. chinensis* to *R. lucieae* than *R. rugosa* (Wissemann and Ritz, 2005; Fougère-Danezan et al., 2015; Debray et al., 2019; Jeon et al., 2024), more inter-chromosomal collinear blocks were identified between *R. chinensis* and *R. lucieae*, which also prevailed in the genomic study of *R. lucieae* ‘Basye’s thornless’ (Zhong et al., 2021), compared to between *R. chinensis* and *R. rugosa* (Fig. 2C; Supplementary Fig. S3). Given the intra-genomic collinear blocks and increased gene content within the genome of *R. lucieae* (Fig. 1; Table 1), putative gene duplication may have influenced the genomic evolution of *R. lucieae*, which aligned with the pronounced expansion of gene families in *R. lucieae* lineage detected from the gene family evolution analysis (Fig. 2C).

**Fig. 2.**
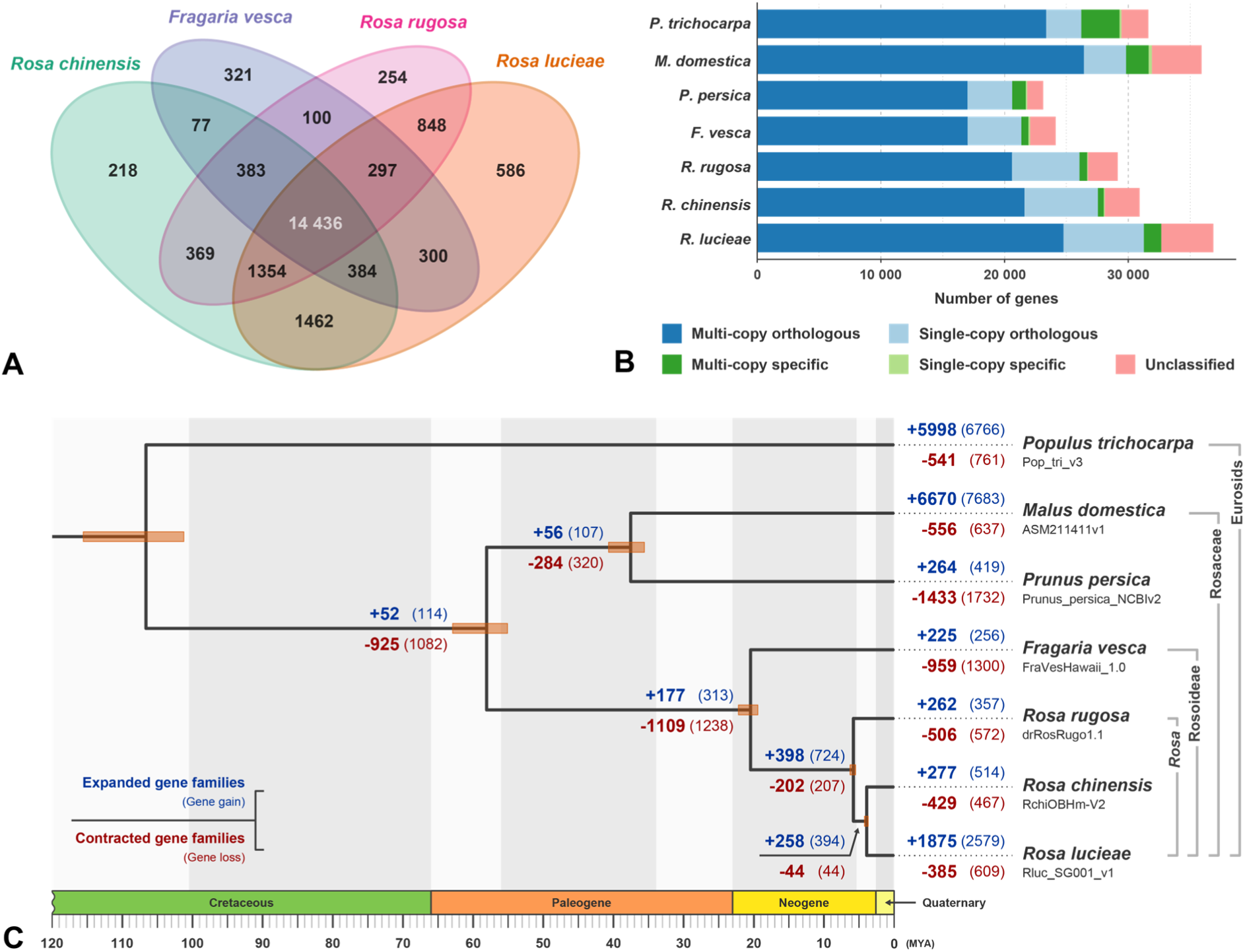
Comparative genomic and evolutionary analyses of *Rosa lucieae*. **A)** Venn diagram of orthologous gene families among *Rosa lucieae*, *R. chinensis*, *R. rugosa* and *Fragaria vesca*. **B)** Gene family classification profile of protein-coding genes of *R. lucieae* and other close reference species. **C)** Chronogram and gene family evolution of *R. lucieae* and other close reference species. In the Bayesian phylogenetic tree, the posterior probability for every branch was 1.0.

### Rapid evolution of salt stress-related gene families in Rosa lucieae

The ortholog group inference analysis identified 2228 single-copy orthologous gene families shared among the genomes of *R. lucieae*, five other rosaceous species, and the outgroup taxa *Populus trichocarpa*. The Bayesian phylogenetic inference using the single-copy orthologous gene set corroborated the taxonomic classification of the species, resolving robust monophyly of the family Rosaceae, subfamilies Rosoideae and Amygdaloideae, and genus *Rosa* respectively (Fig. 2C; Supplementary Table S6). Within the genus *Rosa*, the closer relationship between *R. lucieae* and *R. chinensis* was inferred compared to with *R. rugosa*, which is congruent with the previous phylogenetic inferences of the genus *Rosa* using conventional molecular markers (Wissemann and Ritz, 2005; Fougère-Danezan et al., 2015). In pairwise comparisons of Rosoideae species on distributions of the synonymous substitution rates (Ks) of orthologous genes, orthologous Ks values between *R. lucieae* and *R. chinensis* were lower (mode Ks = 0.021) than those between *R. rugosa* and other *Rosa* species (mode Ks = 0.033–0.035), and between *F. vesca* and *Rosa* species (mode Ks = 0.187– 0.199; Supplementary Fig. S4). This result suggested more recent divergence of *R. lucieae* and *R. chinensis* compared to their divergence with *R. rugosa*, aligning with the divergence time estimation (Fig. 2C).

Based on the inferred orthologous gene families and species phylogeny, the gene family evolution was inferred within each phylogenetic lineage. While contractions of gene families were predominantly involved in the evolution of most other rosaceous lineages, the evolution of *R. lucieae* involved the pronounced expansion of gene families, from the stem lineage of genus *Rosa* (Fig. 2C). After the divergence with *R. chinensis*, 1875 gene families were expanded with 2579 more genes, whereas 385 gene families were contracted with 609 less genes within the lineage of *R. lucieae*. Among the expanded or contracted gene families involved in the *R. lucieae* lineage, 283 gene families were inferred to be rapidly evolving in this lineage. For the *R. lucieae* genes of rapidly evolving families, GOs related to plant signaling, secondary metabolites, salinity stress, root system, homeostasis, calcium ion, protein repair, or defense mechanism were significantly enriched (Fig. 3; Supplementary Table S7). Likewise, based on their KEGG orthology (KO) annotations, KEGG pathways related to the steroid hormones including BRs, secondary metabolites, and stress responses were significantly enriched (Supplementary Fig. S5).

**Fig. 3.**
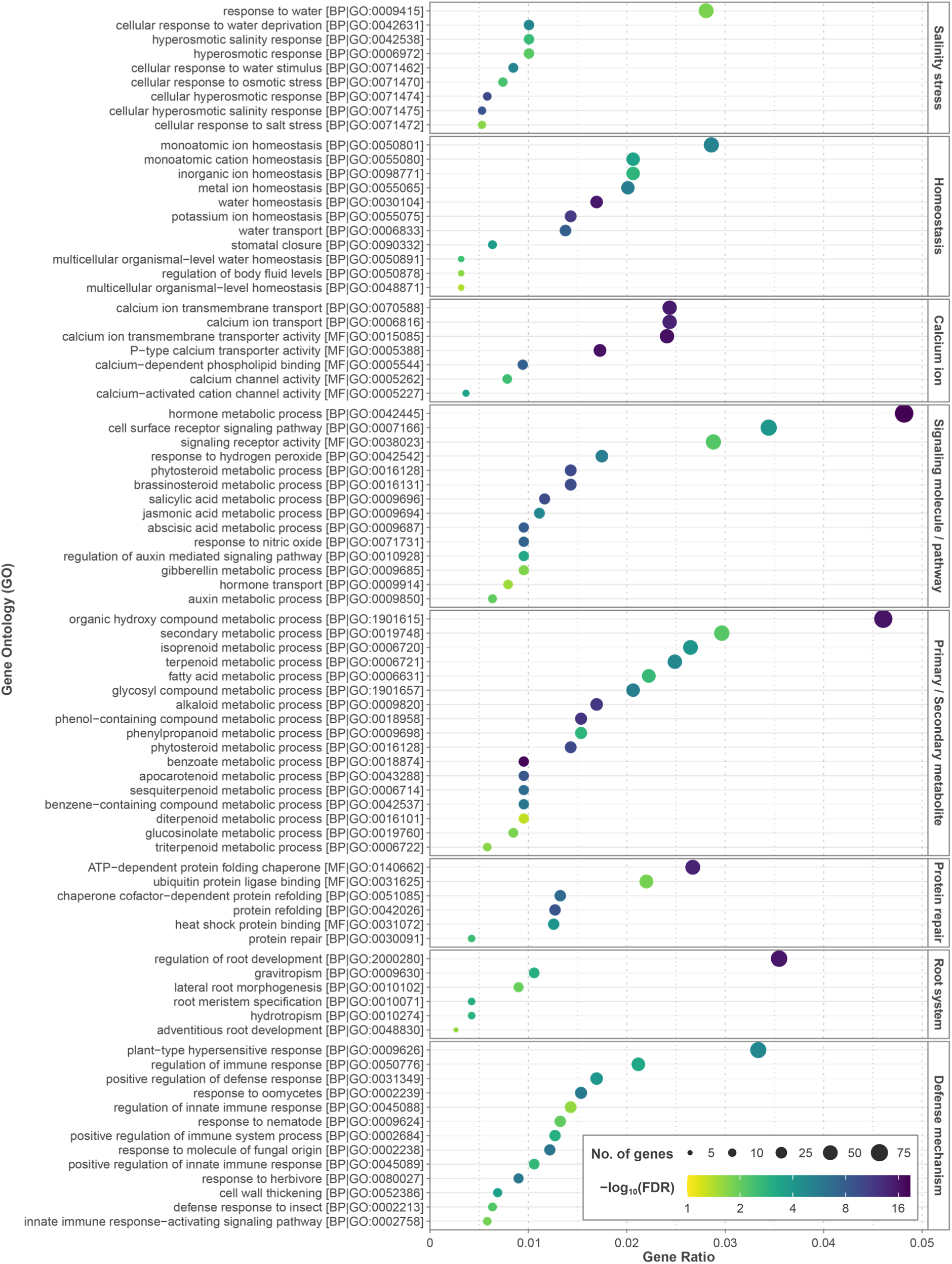
Gene ontology (GO) enrichment analysis on *Rosa lucieae* genes within the rapidly evolving gene families based on the gene family evolution analysis. BP: biological process; MF: molecular function; CC: cellular component; FDR: false discovery rate.

Within the genus *Rosa*, paralogous Ks values were greater (mode Ks = 0.109–0.122) than Ks values of orthologous genes, and their distributions were mostly overlapped with each other (Fig. 4A). This implied that paralogous genes in most gene families were likely duplicated in their common ancestral lineage after the divergence with *F. vesca*, while the orthologous genes have been accumulating unique genetic variation after their divergence by duplications. Particularly with paralogous genes annotated with the GO term ‘response to salt stress’ (GO:0009651) or its child terms, Ks values in *R. lucieae* were remarkably decreased (mode Ks = 0.017) compared to Ks values of all paralogous genes (mode Ks = 0.114), whereas Ks values in other *Rosa* species and *F. vesca* were barely changed or slightly increased (mode Ks = 0.102–0.169; Fig. 4B). The unique Ks distribution of paralogous salt stress-related genes in *R. lucieae* with lower values indicated the recent duplications and divergence of these genes specifically within the evolution of *R. lucieae*.

**Fig. 4.**
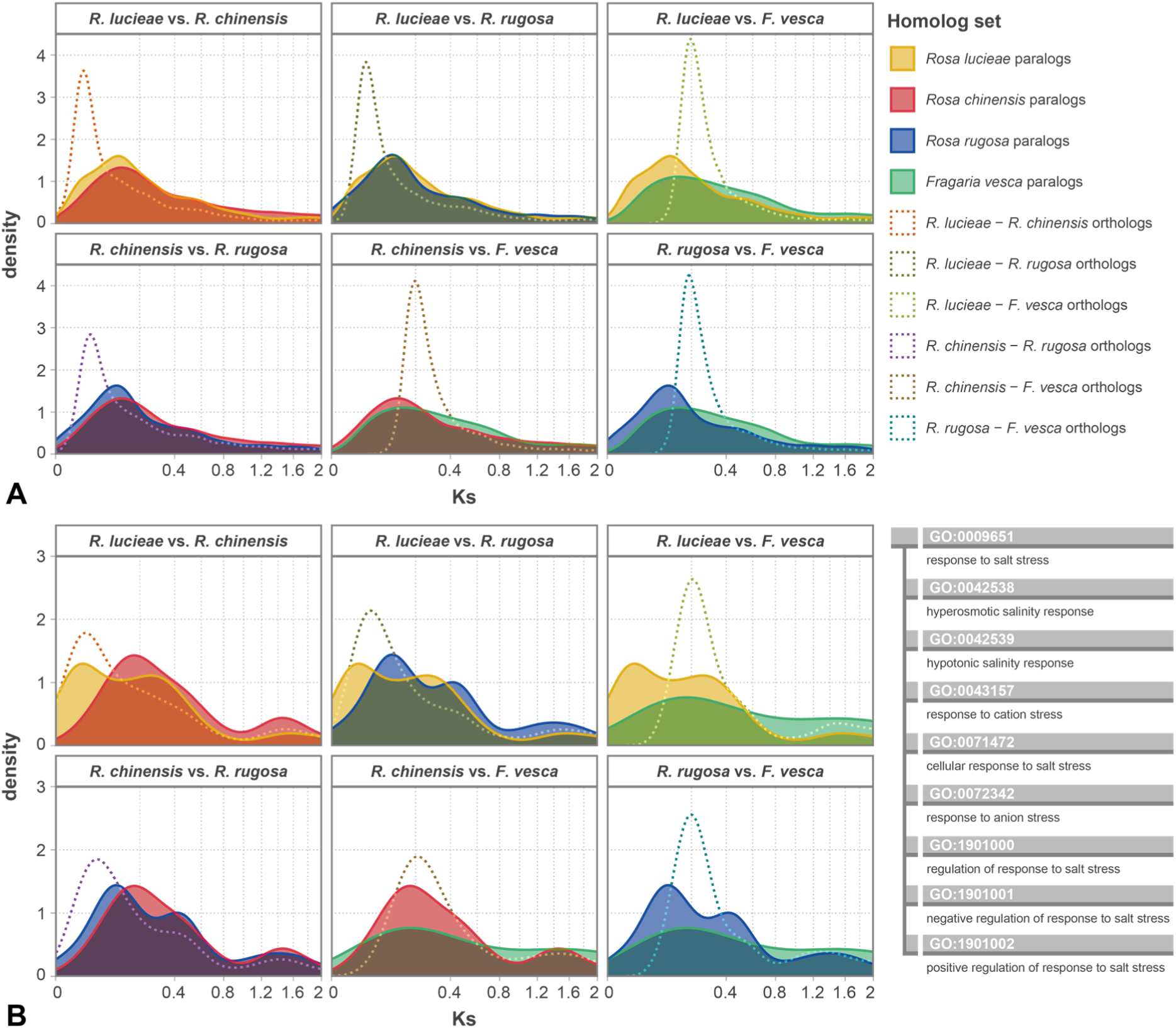
Density plots of the synonymous substitution rates (Ks) of pairs of homologous genes within the four Rosoideae species. **A)** Pairwise comparisons of Ks distribution of paralogous and orthologous genes within pairs of the Rosoideae species. **B)** Pairwise comparisons of Ks distribution of paralogous and orthologous genes annotated with the gene ontology (GO) term ‘response to salt stress’ (GO:0009651) or its child terms (left), of which the ancestor chart is shown on the right side.

### Duplication and divergence of salt tolerance-related genes in Rosa lucieae

Within the rapidly evolving gene families of *R. lucieae*, GOs related to responses to salt and osmotic stresses were significantly enriched (Fig. 3; Supplementary Table S7), suggesting possible influence of gene duplications or divergence of *R. lucieae* on adaptation to salt stress. Moreover, the unique Ks distribution with lower values of paralogous *R. lucieae* genes annotated with GOs related to responses to salt stress corroborated their recent divergence compared to those genes in other close species or other genes in *R. lucieae* (Fig. 4). Given that gene duplication and copy-number variation can contribute to adaptive evolution by increasing the genetic diversity or product dosage (Ohno, 1970; Kondrashov, 2012; Magadum et al., 2013), these results substantiated the influence of duplications and divergence of salinity response-related genes on the unique adaptation to salinity in *R. lucieae*. Similarly, the duplications of various salinity response-related genes were identified in the evolution of salt tolerance of the saltwater cress, *Eutrema salsugineum* (syn. *Thellungiella salsuginea*), which is a close halophytic relative of *Arabidopsis thaliana* (Wu et al., 2012; Monihan et al., 2019).

In addition to GOs related to responses to salt and osmotic stresses, GOs related to ABA signaling, Ca^2+^ signaling and ROS signaling, and KEGG pathways related to MAPK signaling were also significantly enriched in the rapidly evolving gene families of *R. lucieae*, which are principal signaling pathways of salinity response in plants (Fig. 3; Supplementary Fig. S5; Supplementary Table S7; Zhu, 2001; Botella et al., 2005; Van Zelm et al., 2020). The genetic evolution related to early signaling under salt stress possibly influenced the adaptation to salinity of *R. lucieae* by enabling a rapid and/or enhanced response to salt stress. Within the genome of *R. lucieae*, the duplications and divergence of various salt stress signaling-related genes were identified, including genes of Ca^2+^ channels such as cyclic nucleotide-gated ion channel 1 (CNGC1) (Table 2; Supplementary Figs. S6 and S7; Supplementary Table S8).

**Table 2.**
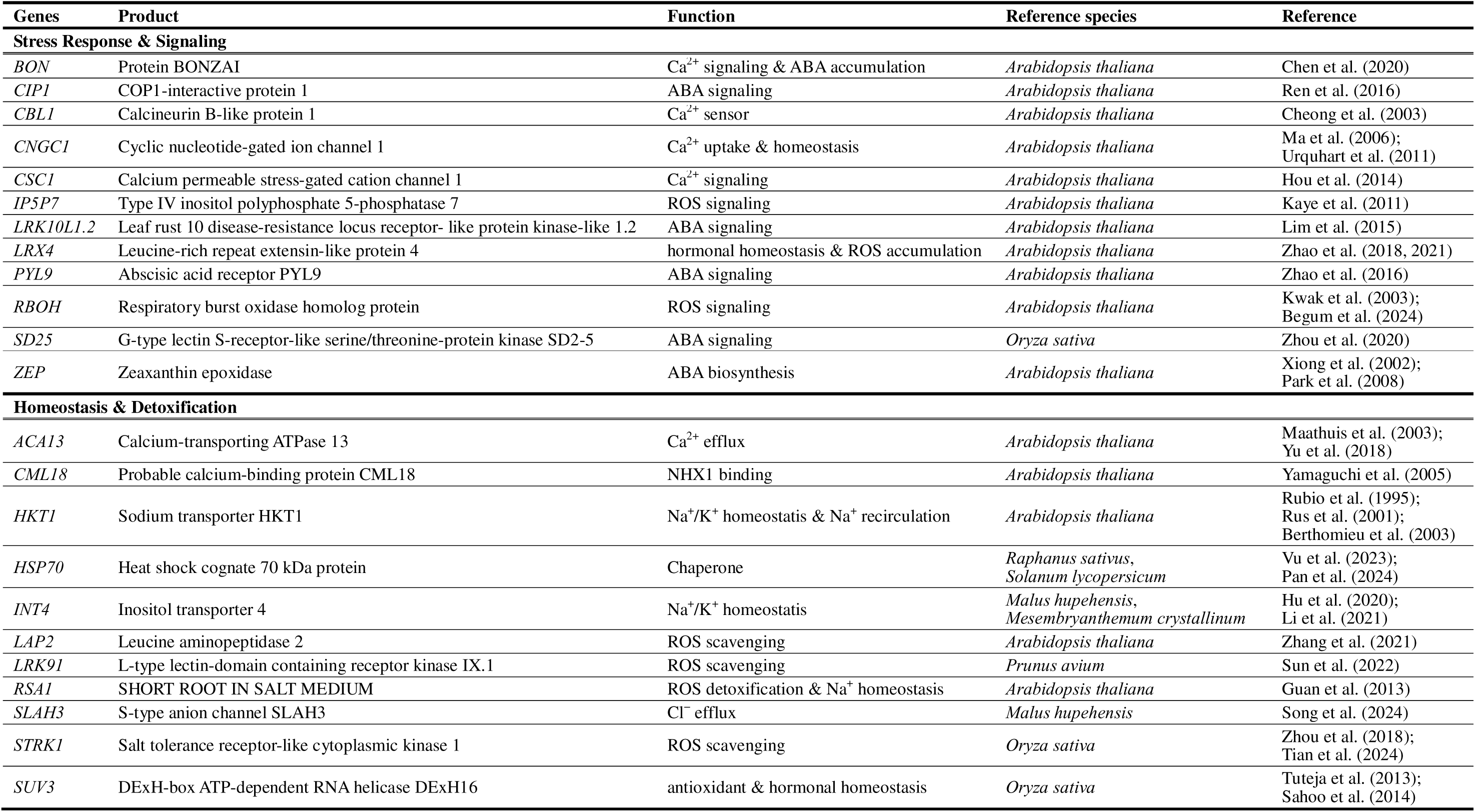
List of candidates of salt tolerance-related genes duplicated in *Rosa lucieae*.

CNGC1 is a member of cyclic nucleotide-gated channels (CNGCs), which are Ca^2+^- permeable cation channels involved in signal transduction related to stress tolerance in plants (Sanders et al., 2002; Jha et al., 2016). While some other members of CNGCs in *A. thaliana*, such as AtCNGC19 and AtCNGC20, are known to be involved in salt stress (Kugler et al., 2009), the function of *Arabidopsis* CNGC1 in stress tolerance is less understood but Pb^2+^ tolerance (Sunkar et al., 2000). Even though the direct relationship of AtCNGC1 to salt stress is barely known, several recent studies suggested the role of AtCNGC1 in Ca^2+^ uptakes at roots, accumulation of Ca^2+^, and maintaining proper Ca^2+^ levels in plants (Ma et al., 2006; Urquhart et al., 2011). In the genome of *R. lucieae*, the copy number of CNGC1 genes was increased compared to other close Rosoideae species (Supplementary Fig. S6). Gene duplication of *R. lucieae CNGC1* (*RlCNGC1*) was not likely to originate from a specific homologous gene cluster, but has occurred in diverse homologous gene clusters independently (Supplementary Fig. S6A). The synteny of orthologous *CNGC1* mostly conflicted with the synteny of genomic collinear blocks, and several copies of *RlCNGC1* were inserted in multiple unplaced contigs of *R. lucieae* (Supplementary Fig. S6B, C).

The increased number of *RlCNGC1* may have affected the accumulation of Ca^2+^ in *R. lucieae* by regulating Ca^2+^ uptake at roots, which can induce a rapid increase in Ca^2+^ levels after salinity stress (Ma et al., 2006; Kader and Lindberg, 2010). *Rosa lucieae* can be more Ca^2+^-sensitive than other roses with the increased gene copy numbers of Ca^2+^ sensors such as Calcineurin B-like protein 1 (CBL1), of which overexpression in transgenic *A. thaliana* plants enhanced salt tolerance (Albrecht et al., 2003; Cheong et al., 2003). After the elevation of Ca^2+^ levels induced by salt stress, the increased gene copy numbers of salt stress-related signaling proteins including ion channels, receptors and kinases possibly influenced the subsequent ABA, Ca^2+^ and ROS signaling in *R. lucieae* (Table 2). For example, transgenic *A. thaliana* plants overexpressing an ABA receptor, pyrabactin resistance 1-like 9 (PYL9), exhibited improved osmoregulation (Zhao et al., 2016). Likewise, transgenic *A. thaliana* plants overexpressing zeaxanthin epoxidase (ZEP), which is pivotal in ABA biosynthesis, also exhibited enhanced salt tolerance (Park et al., 2008). Thus, overexpression of stress response- and signaling-related genes by duplication may have contributed to the adaptive evolution of *R. lucieae* in saline environments. Given the increased genetic diversity after duplication (Ohno, 1970; Kondrashov, 2012), the duplicated gene pairs were also possibly led to asymmetric evolution, resulting in functional and conditional differences in stress signaling (VanderSluis et al., 2010).

Because the early responses of salt stress including generation of ABA and ROS can damage and impair plant systems, the downstream detoxification and recovery are another pivotal mechanism in salt tolerance (Van Zelm et al., 2020). The significant enrichment of metabolite-, ionic and osmotic homeostasis-, and protein repair-related GOs, and secondary metabolite- and protein processing-related KEGG pathways implied the putative evolution of ion- or ROS-detoxification strategies in *R. lucieae* (Fig. 3; Supplementary Fig. S5; Supplementary Table S7). Within the genome of *R. lucieae*, the duplications and divergence of various detoxification- and homeostasis-related genes were also identified, including genes of ion transporters such as auto-inhibited Ca^2+^-ATPases (ACAs) (Fig. 5; Table 2; Supplementary Fig. S7; Supplementary Table S8).

**Fig. 5.**
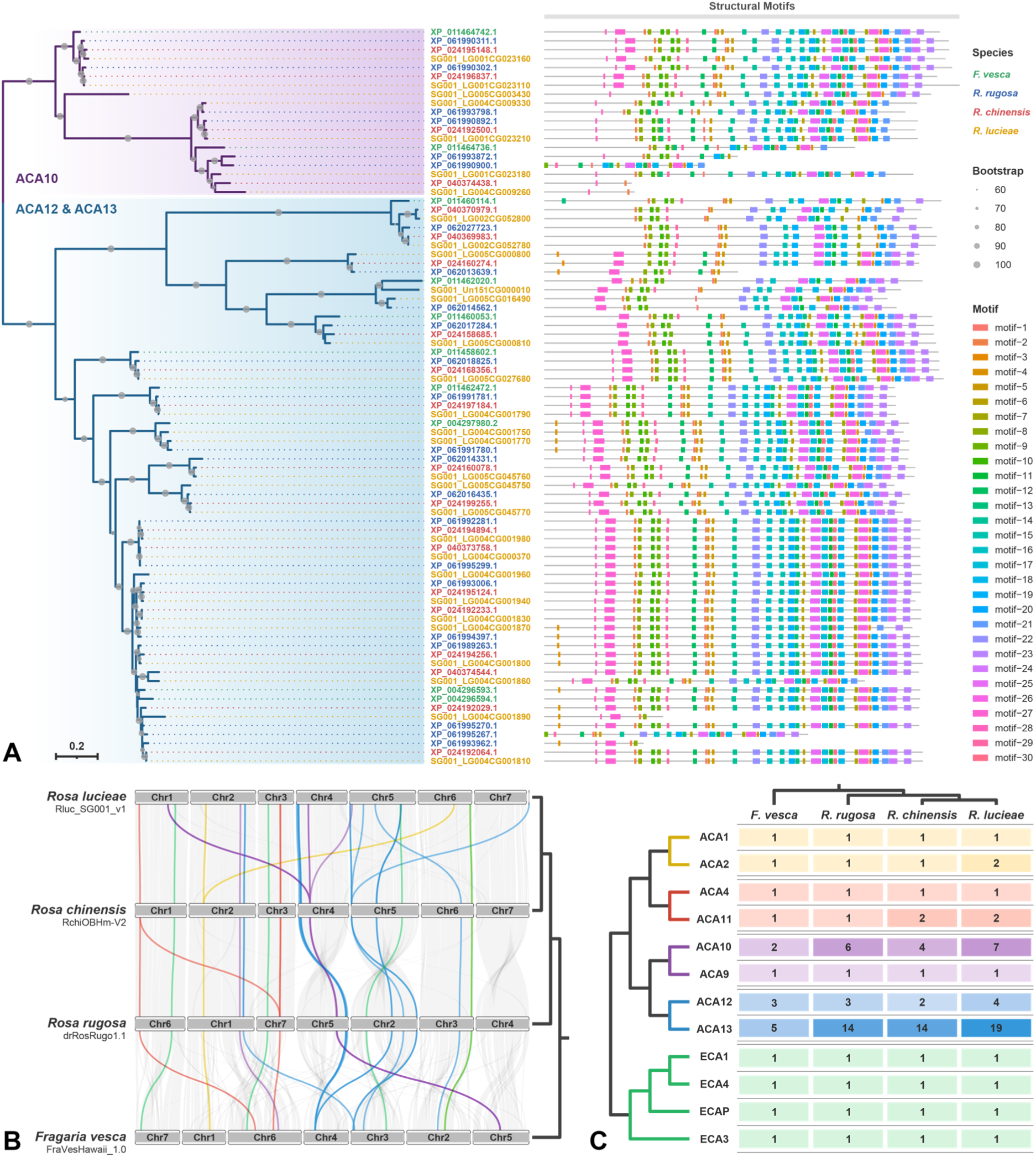
Duplications and divergence of ACA genes within Rosoideae. **A)** Unrooted gene tree of ACA10, ACA12 and ACA13 genes (left) and the structural motifs in each ACA gene (right). **B)** Ideogram of the four Rosoideae species showing synteny of ACA and ECA genes between species. Colored lines represent the synteny relationships of orthologous ACA and ECA gene pairs, and gray lines represent the synteny relationships of genomic collinear blocks. The colors of lines represent the clusters within the ACA and ECA gene family (see subfigure C). The cladogram on the right side represents the species relationships. **C)** Copy number variation of ACA and ECA genes within the genomes of four Rosoideae species. The numbers in blocks and the color shades of blocks represent the gene copy numbers.

ACAs and endoplasmic reticulum-type Ca^2+^-ATPases (ECAs) are P-type Ca^2+^-ATPases found in plants, of which a primary role is terminating Ca^2+^-mediated signaling by Ca^2+^ efflux from the cytosol (Bose et al., 2011; Park and Shin, 2022). Thus, ACAs and ECAs are pivotal in the removal of excessive Ca^2+^ and restoration after stress responses. ACAs are generally classified into four conserved clusters in flowering plants: L) ACA1, ACA2 and ACA7 in cluster 1, L) ACA4 and ACA11 in cluster2, L) ACA12 and ACA13 in cluster 3, and L) ACA8, ACA9 and ACA10 in cluster 4 (Yu et al., 2018). The unique gene functions of each cluster in stress responses were investigated in several model plants including *A. thaliana* and *Oryza sativa* (Bose et al., 2011; Huda et al., 2013; Yu et al., 2018; Park and Shin, 2022). Within the genome of *R. lucieae*, genes of various members of ACA family were duplicated, including genes of ACA10, ACA12 and ACA13 (Fig. 5). Similar to *RlCNGC1*, the duplications of *R. lucieae ACA10*, *ACA12* and *ACA13* have occurred in diverse homologous gene clusters independently (Fig. 5A, B). The copy numbers of *ACA13* of *Rosa* species were notably increased compared to their relative *F. vesca*, and the copy number of *ACA13* of *R. lucieae* was increased again within the genus *Rosa* (Fig. 5A, C).

The excessive Na^+^, Cl^−^ and Ca^2+^ resulting from high salinity can cause the ion toxicity by themselves, and also induce severe stress responses such as overaccumulation of ROS (Zhu, 2001; Munns and Tester, 2008). In addition to *RlACA13*, the duplications and divergence of *R. lucieae* genes of calmodulin-like protein 18 (CML18), high-affinity K^+^ transporter 1 (HKT1), inositol transporter 4 (INT4), SHORT ROOT IN SALT MEDIUM 1 (RSA1), and slow anion channel 1 homolog 3 (SLAH3) have possibly affected alleviation and detoxification of excessive Na^+^, Cl^−^ and Ca^2+^, rescuing *R. lucieae* from salt stress. For examples, the point mutation in *A. thaliana HKT1* increased the Na^+^ tolerance with reduced Na^+^ uptake (Rubio et al., 1995), and three paralogous copies of *MhINT4* in the Chinese crab apple (*Malus hupehensis*; Rosaceae) were differently involved in salt tolerance, in which *MhINT4.2* was particularly up-regulated under salt stress (Hu et al., 2020). Furthermore, transgenic *M. hupehensis* plants overexpressing SLAH3 exhibited increased Cl^−^ efflux rate and reduced ROS accumulation after ABA signaling. Likewise, transgenic *A. thaliana* plants overexpressing RSA1 exhibited enhanced expression of salt overly sensitive 1 (SOS1), a Na^+^/H^+^ antiporter crucial for Na^+^ homeostasis under salt stress, and genes encoding oxidoreductase and peroxidase, resulting in Na^+^ and ROS detoxification (Guan et al., 2013). In addition to *RlRSA1* and *RlSLAH3*, the duplications and divergence of *R. lucieae* genes of leucine aminopeptidase2 (LAP2), L-type lectin-domain containing receptor kinase IX.1 (LRK91), salt tolerance receptor-like cytoplasmic kinase 1 (STRK1), and suppressor of Var 3 (SUV3) are possibly responsible for scavenging and detoxification of ABA-induced ROS. The various receptor-like kinases such as LRK91 and STRK1 are well known to be involved in responses to plant abiotic stresses (Ye et al., 2017; Gandhi and Oelmüller, 2023). For instance, transgenic sweet cherry (*Prunus avium*; Rosaceae) plants overexpressing LRK91 homolog (PaLectinL16) and transgenic *O. sativa* plants over-expressing STRK1 exhibited enhanced antioxidant ability under salt stress (Zhou et al., 2018; Sun et al., 2022). Likewise, transgenic *O. sativa* plants overexpressing SUV3 also exhibited reduced ROS production with increased antioxidant activities, and increased growth rates with higher content of phytohormones under salt stress (Tuteja et al., 2013; Sahoo et al., 2014). Furthermore, given the diverse roles of 70 kD heat shock proteins (HSP70s) as molecular chaperones in abiotic stress tolerance, the increased copy number and genetic diversity of *RlHSP70* possibly influenced various abiotic stress tolerance in *R. lucieae*, including salt tolerance (Vu et al., 2023; Pan et al., 2024). Thus, increased gene copy numbers and diversity after duplications of homeostasis- and detoxification-related genes of *R. lucieae* may have contributed to the evolution of increased viability and rapid recovery from severe salt stress, resulting in the vigorous growth of *R. lucieae* in saline environments.

BRs are a group of natural steroid hormones broadly found in plants, which are considered to be related to plant growth, development and stress response. Although their roles in plant growth and development are well established, the mechanism of stress resistance induced by BRs is poorly understood (Krishna, 2003). Nevertheless, several recent studies suggested BRs enhance the stress resistance with the complex interaction with other hormones, including cross-talk between BRs and ABA (Krishna, 2003; Choudhary et al., 2012). Under salt stress, BRs modulate the gene transcription and ROS levels by acting as signaling molecules and interacting other phytohormones (Anwar et al., 2018; Kumar et al., 2022). Although not many BR-related genes affecting salinity resistance are known in plants, pronounced enrichment of BR-related GOs and KEGG pathways suggested the putative influence of the rapid evolution of BR-related genes on salt tolerance of *R. lucieae* (Fig. 3; Supplementary Fig. S5; Supplementary Table S7).

### Modes of gene duplication in adaptive evolution of Rosa lucieae

There are four different modes of gene duplication known so far: tandem duplication, whole-genome duplication, chromosomal segmental duplication, and single gene transposition-duplication (Freeling, 2009). Among those, whole-genome duplication is less likely to affect the gene duplications and divergence in *R. lucieae*, considering its similar genome size and nucleic DNA amount compared to its close relatives (Yokoya et al., 2000). Except for whole-genome duplication, salt tolerant-related genes of *R. lucieae* have been possibly duplicated through any other modes of duplication. Especially, all the three modes of duplication are likely to influence the divergence of *RlCNGC1*. The clusters of adjacent *CNGC1* paralogs located on chromosome 1 of *R. lucieae*, which are conserved in Rosoideae species, may have originated from the ancestral tandem duplication within the Rosoideae lineage (Supplementary Fig. S6B; Supplementary Table S8). One of these *CNGC1* clusters in *R. lucieae* was duplicated into unplaced contig 62, hypothesizing a recent segmental duplication event in *R. lucieae* probably increased the copy number of *RCNGC1* (Supplementary Fig. S6B, C). The close genetic distances of *RlCNGC1* genes between the two clusters on chromosome 1 and unplaced contig 62 corroborated their recent divergence (Supplementary Fig. S6A). On the other hand, many *CNGC1* genes of which synteny was not aligned with the synteny of genomic collinear blocks were dispersed on various loci of whole genomes (Supplementary Fig. S6B). This distribution can stem from transposition-duplication, which relocates genes to new and arbitrary loci and blurs the syntenic evidence (Freeling, 2009). The dense distributions of LTR elements and DNA repeat elements near the duplicated *RlCNGC1* genes on novel positions supported the possible transposition-duplication of *RlCNGC1* (Supplementary Fig. S6C). The various modes of duplication of *RlCNGC1* suggested that all duplication modes but whole-genome duplication may have influenced the recent and rapid adaptive evolution of *R. lucieae* in saline environments.

### Evolution of other stress tolerance in Rosa lucieae

In addition to salt stress in their natural habitat, wild *R. lucieae* plants are exposed to various biotic and abiotic stresses, and become adapted and tolerate to these stresses. The Ca^2+^ and ABA signaling, and ROS accumulation are common processes of responses to diverse biotic and abiotic stresses (Mauch-Mani and Mauch, 2005; Fujita et al., 2006; Boudsocq and Sheen, 2010; Brosché et al., 2010; Rock et al., 2010). Moreover, some genes responsible to salt tolerance are also related to other abiotic stress, especially drought stress, which induces many common responses with salt stress by causing osmotic stress (Bartels and Sunkar, 2005). For example, the overexpression of CBL1 enhanced drought tolerance in transgenic *A. thaliana* plants as well, in addition to salt tolerance (Albrecht et al., 2003; Cheong et al., 2003). Similarly, closely related genes to putative salt tolerance-responsible genes in *R. lucieae* may be related to other stress tolerance of *R. lucieae*. For example, similar to *RlACA13*, the copy numbers of *RlACA10* and *RlACA12* were increased compared to close relatives of *R. lucieae* (Fig. 5C). Despite the genetic structures and function of Ca^2+^ efflux similar to ACA13, ACA10 and ACA12 are responsible to plant immune response and defense to pathogens and insects in *A. thaliana* but not salt tolerance (Frei dit Frey et al., 2012; Yu et al., 2018; Fotouhi et al., 2022). This result suggested that gene duplications and divergence also enhanced stress response, signaling and restoration of *R. lucieae* under various biotic and abiotic stresses.

Aligning with the contribution of *R. lucieae* to the improvement in resistance to diseases and pathogens in rose breeding (Noack, 2003; Byrne et al., 2005), GOs and KEGG pathways related to plant immunity and defense mechanism were significantly enriched in the rapidly evolving gene families of *R. lucieae* (Fig. 3; Supplementary Fig. S5; Supplementary Table S7). Likewise, the significant enrichment of GOs and KEGG pathways related to phytohormones and secondary metabolites responsible for plant immunity and defense supported the enhanced resistance to biotic stresses in *R. lucieae*. The phytohormones salicylic acid (SA), JA and ethylene are renowned for their roles in biotic stress responses (Fujita et al., 2006), and GOs related to all these hormones were significantly enriched in the rapidly evolving gene families of *R. lucieae* (Fig. 3; Supplementary Table S7). Given that SA and ethylene can interact with BRs to enhance salt tolerance (Divi et al., 2010; Choudhary et al., 2012), the evolution of salt tolerance may be closely associated with the evolution of enhanced defense to pathogens and insects in *R. lucieae*. The secondary metabolites including terpenoids, phenolics, alkaloids and flavonoids are pivotal for plants in defense and adaptation to environmental stresses (Akula and Ravishankar, 2011; Erb and Kliebenstein, 2020). Particularly, the importance of secondary metabolites in salinity tolerance in plants was highlighted with their crucial role as antioxidants alleviating oxidative stress induced by high salinity (Isah, 2019). The recent multi-omics study also emphasized the importance of flavonoid and flavonol metabolism in response to salt stress in roses (Ren et al., 2024).

Within secondary metabolites, plant volatiles particularly act on attracting pollinators as floral scent components or deterring herbivory as insect repellents, and non-volatiles are involved in various defense mechanism distinct from volatiles (Pichersky and Gershenzon, 2002; Unsicker et al., 2009; Tholl, 2015). The earlier comparative genomic investigations of rose species suggested that the duplication and divergence of volatile terpenoid metabolism-involved Nudix hydrolase 1 (NUDX1) genes modulate the content of scent compounds of rose flowers (Conart et al., 2022; Shang et al., 2024). However, the copy numbers of *NUDX1* were not only decreased in *R. lucieae* (four copies in *R. lucieae*, nine in *R. chinensis*, and 15 in *R. rugosa*), but the amount of volatile compounds was also reduced in flower petals of *R. lucieae* (Feng et al., 2022; Shang et al., 2024). Moreover, different profiles of paralogous *NUDX1* in involvement in floral scent were identified between *R. chinensis* and hybrid cultivars of *R. lucieae* (Conart et al., 2023). In rose flowers, the chemical composition of volatile compounds was notably different among rose species, and the interspecific relationships based on characteristics of floral scent were inconsistent with the species phylogeny (Debray et al., 2019; Feng et al., 2022; Noh et al., 2024; Jeon et al., 2024; Shang et al., 2024), which implied the independent and unique evolution of secondary metabolites in different rose species. This diversity and complexity of secondary metabolites among closely related species can be easily affected by environmental factors (Borghi et al. 2019; Paul et al., 2022). Moreover, the biosynthetic pathways of volatiles are also responsible for production of various non-volatile secondary metabolites involved in stress resistance (Paul et al., 2022). Given the enhanced resistance to biotic and abiotic stresses in *R. lucieae*, the unique biosynthetic mechanism of secondary metabolites in *R. lucieae* may have alternatively evolved to complement resistance to environmental stresses. For example, the prominently enriched GOs of saponin and glycoside metabolism in *R. lucieae* may be responsible for defense to herbivory, diseases and pathogens (Supplementary Table S7; Tholl, 2015). Further biochemical investigation of *R. lucieae* will contribute to understanding protective compounds of roses and their mechanisms of defense to diseases, pathogens and other environmental stresses, promising our understanding of hardiness in rose breeding programs.

## DISCUSSION

*Rosa lucieae* inhabits open and exposed rocky slopes or banks, especially along the sea shores and riverside at low elevation (Rehder, 1940; Ohba, 1993; Ku and Robertson, 2003), where the salts are constantly accumulated in their soils. In these natural habitats, *R. lucieae* has evolved to adapt to saline environments, resisting to and growing vigorously under salt stress. Although *R. rugosa* is another rose species renowned for salt tolerance, its earlier genomic study could not find any difference in genetic component related to salt tolerance, but only find the duplications of two putative genes (*DRIP2* and *PTR3*) related to water deprivation (Chen et al., 2021). However, the first chromosome-level genome of wild *R. lucieae* from the present study shed light on the adaptive evolution to salt stress in roses.

Utilizing high-throughput genome sequencing techniques, the genome of wild *R. lucieae* was sequenced and assembled into 544.7 Mb-long chromosome-level genomic sequences. This assembly presents the genomic basis of a wild rose, which provides insights into the nature of roses. The chromosome-level genome assembly of *R. lucieae* indicated the conserved genomic structure with other comprehensive rose genomic assembly of *R. chinensis* and *R. rugosa* (Supplementary Fig. S3). Nevertheless, coupled with the increased genome size, the number of protein-coding genes within the genome of *R. lucieae* was increased with gene duplications (Table 1). This remarkable gene duplication was identified as a major driving force behind the adaptive evolution in salt tolerance of *R. lucieae*. The diverse genes responsible for various processes of salt tolerance in plants were duplicated in *R. lucieae*, including Ca^2+^ (*BON*, *CBL1*, *CNGC1* and *CSC1*), abscisic acid (*BON*, *CIP1*, *LRK10L1.2*, *PYL9*, *SD25* and *ZEP*), and ROS signaling (*IP5P7*, *LRX4* and *RBOH*), ion homeostasis (*ACA13*, *CML18*, *HKT1*, *INT4*, *RSA1* and *SLAH3*), ROS detoxification (*LAP2*, *LRK91*, *SLAH3*, *STRK1* and *SUV3*), protein repair (*HSP70*), and growth recovery (*LRX4* and *SUV3*) (Table 2; Supplementary Fig. S7). Moreover, the rapid evolution of gene families related to phytohormones including BRs, SA and ethylene also possibly enhanced tolerance to various stresses including salinity. The unique evolution in composition of secondary metabolites may have allowed the adaptation to environmental stresses and defense to diseases and pathogens. These results substantiate an important role of gene duplication in the adaptive evolution of plants, and shed light on the nature of adaptive evolution of *R. lucieae* in saline environments. The genomic basis of salt tolerance of *R. lucieae* will pave the way for the advancement in rose breeding programs by contributing to enhanced stress resistance in rose cultivation. Particularly, understanding genomic mechanism of resistance to environmental stresses in roses and its utilization will significantly contribute to future horticultural and agricultural industries with recovered or enhanced productivity and quality of rose flowers or other rosaceous crops in ongoing global urbanization and climate changes. A further comparative analysis of expression profiles in *R. lucieae* will corroborate influence of the putative salt tolerance-responsible genes on the adaptation of *R. lucieae* to saline environments. Additionally, further genome-wide association studies and comparative genomic studies including diverse *R. lucieae* accessions from various sources will unveil the influence of genetic mutations affecting salt tolerance in roses.

## MATERIALS AND METHODS

### Sample collection and preparation

To develop a genomic sequence of *R. lucieae*, a wild individual of *R. lucieae* (accession ID: SG001; NCBI BioSample accession: SAMN39614440) was collected from a coastal field in Jeju, Korea, and its voucher specimen was deposited into the Ha Eun Herbarium (SKK) of Sungkyunkwan University. Genomic DNA (gDNA) was isolated from fresh young leaves of the accession using the Exgene Plant SV mini kit (GeneAll Biotechnology Co., Ltd., Seoul, Korea), following the manufacturer’s instructions. For the quality control, the isolated gDNA was quantified and assessed using QuantiFluor dsDNA System (Promega Corp., WI, USA) and the 2100 Bioanalyzer System (Agilent Technologies, Inc., CA, USA). Additionally, flower buds (BioSample accession: SAMN40548981) and young leaves (BioSample accession: SAMN40548982) were collected from the same individual to provide evidence for the splice site prediction. The samples were stored in RNAlater solution (Thermo Fisher Scientific Inc., MA, USA) immediately after the collection to stabilize their RNAs. Total RNAs were isolated using Maxwell RSC Plant RNA Kit and Instrument (Promega) by Macrogen, Inc. (Seoul, Korea). For the quality control, the isolated RNAs were quantified and assessed using Quant-iT RiboGreen RNA Assay Kit (Thermo Fisher Scientific) and the 2100 Bioanalyzer System.

### Molecular sequencing and genome assembly

The gDNA was prepared into respective libraries for Illumina HiSeq short-read and PacBio SMRT long-read sequencing platforms. The short-read sequencing library was prepared by random fragmentation, followed by 5′- and 3′-adapter ligations, using the TruSeq Nano DNA kit (Illumina, Inc., CA, USA). The fragmented library was sequenced via Illumina HiSeq platform (Illumina) by Macrogen after the size selection. The PacBio SMRTbell library was prepared using SMRTbell Express Template Prep Kit (Pacific Biosciences of California, Inc., CA, USA), following the manufacturer’s instructions. The large fragments longer than 20 kb were selected using BluePippin system (Sage Science, Inc., MA, USA), and the large-insert library was sequenced via PacBio Sequel platform (Pacific Biosciences) by DNALink, Inc. (Seoul, Korea). The total RNAs were prepared into sequencing libraries by purifying poly-A containing mRNAs, random fragmentation, cDNA synthesis, and 5′- and 3′-adapter ligations, using the TruSeq Stranded mRNA kit (Illumina). The fragmented libraries were sequenced via Illumina NovaSeq platform (Illumina) by Macrogen.

The genome size, heterozygosity and repeat content of *R. lucieae* were estimated with *k*-mer analysis using Jellyfish v2.1.3 (Marçais and Kingsford, 2011) and GenomeScope v2.0 (Ranallo-Benavidez et al., 2020), after preprocessing short reads by trimming the sequencing adapters, PCR dimers, and low-quality sequences using BBDuk v38.26 (Bushnell, 2014) and removing contaminant reads by mapping reads to NCBI RefSeq complete bacterial, viral and organellar genomes using bowtie2 v2.3.4 (Langmead and Salzberg, 2012). PacBio subreads were error-corrected into consensus reads using SMRT Link v2.3 (Pacific Biosciences), and preprocessed short reads were mapped to the consensus reads to remove contaminant reads using bwa v0.7 (Li, 2013). The preprocessed PacBio consensus reads were *de novo* assembled into primary contigs using a diploid-aware assembler FALCON, and the primary contigs were phased and updated using a diploid assembler FALCON-Unzip (Chin et al., 2016). The phased primary contigs of the diploid genome assembly were curated to remove haplotigs to complete the draft genome assembly using Purge haplotigs v1.1.1 (Roach et al., 2018). Error sequences in the draft genome assembly were corrected using Pilon v1.23, after mapping preprocessed short reads to the draft genome assembly using bwa aligner (Li, 2013; Walker et al., 2014). The *de novo* genome assembly was assessed for the completeness based on the universal single-copy orthologs using BUSCO v5.3.0 and eudicots_odb10, embryophyta_odb10, and viridiplantae_odb10 datasets (Manni et al., 2021). The *de novo* genome assembly was scaffolded into chromosomes using RagTag v2.1.0 (Alonge et al., 2022), referenced from the chromosome-level assembly of *R. lucieae* ‘Basye’s Thornless’ (NCBI GenBank accession: GCA_024704745.1; Zhong et al., 2021). The scaffolded chromosomes were assessed with the genome collinearity analysis with the reference assembly using MUMmer4 v4.0.0rc1 (Marçais et al., 2018), examining any genomic gaps or putative misassembled contigs.

### Genome annotation

The repeat sequences of genome of *R. lucieae* were *de novo* predicted and developed into the *de novo* repeat library using RepeatModeler v1.0.8 (Smit and Hubley, 2008), incorporating complementary repeat annotation methods like RECON v1.08 (Bao and Eddy, 2002), RepeatScout v1.0.5 (Price et al., 2005), and TRF v4.04 (Benson, 1999). The repeat regions of draft genome were identified, classified into subclasses based on the Dfam database v2.0 (Hubley et al., 2016) and RepBase database v23.10 (Bao et al., 2015), and masked using RepeatMasker v4.0.7 (Smit et al., 2013).

Within the genome of *R. lucieae*, protein-coding genes were predicted and annotated by integrating *ab initio*, RNA-seq evidenced, and homology-based gene predictions. For *ab initio* gene prediction, the protein-coding gene models of rosaceous reference genomes of *R. chinensis* ‘Old Blush’ (RefSeq accession: GCF_002994745.2; Raymond et al., 2018), *R. rugosa* (GCF_958449725.1; Ruhsam et al., 2024), *F. vesca* (GCF_000184155.1; Shulaev et al., 2011), *Prunus persica* ‘Lovell’ (GCF_000346465.2; IPGI, 2013), and *Malus domestica* ‘Golden Delicious’ (GCF_002114115.1; Daccord et al., 2017) were trained and optimized respectively using AUGUSTUS v3.1.0 (Stanke and Waack, 2003). With meta-parameters of rosaceous reference gene models, the *ab initio* gene model of *R. lucieae* was predicted within the draft genome using AUGUSTUS v3.1 (Stanke and Waack, 2003). For homology-based prediction, protein-coding genes from reference genomes (Supplementary Table S6) were aligned to the draft genome of *R. lucieae* to identify homologous regions using Exonerate v2.4.0 (Slater and Birney, 2005). For RNA-seq evidenced prediction, each RNA-seq accession was aligned to the draft genome of *R. lucieae* using STAR v2.5.1a (Dobin et al., 2013). Leveraging the RNA-seq alignment, reference gene models of *R. chinensis*, *R. rugosa*, and *F. vesca*, and evidence from *ab initio* and homology-based gene predictions, an integrated consensus gene model was predicted using GeMoMa v1.9 (Keilwagen et al., 2018) based on conserved coding-sequence and intron positions.

A set of protein-coding genes of *R. lucieae* were annotated in their functions. The GOs (Ashburner et al., 2000) and definitions were annotated using blastp v2.15.0 (Camacho et al., 2009), Blast2GO v6.0.3 (Conesa et al., 2005), InterProScan v5.52 (Jones et al., 2014), and Trinotate v4.0.2 (Bryant et al., 2017), based on public databases including NCBI non-redundant protein sequence (nr; accessed on 12 Feb 2024), Gene Ontology Annotation (GOA; v2023.08; Camon et al., 2004), Pfam (v36.0; El-Gebali et al., 2019), KEGG (Kanehisa et al., 2012), UniProtKB/Swiss-Prot (v2024_01; Boutet et al., 2007), and EggNOG (v5.0; Huerta-Cepas et al., 2019).

### Comparative genomic and evolutionary analyses

To infer the genetic evolution of *R. lucieae*, the orthologous gene groups were identified from the gene models of *R. lucieae* and other reference species including *R. chinensis*, *R. rugosa*, *F. vesca*, *P. persica*, *M. domestica*, and *P. trichocarpa* (RefSeq accession: GCF_000002775.4; Tuskan et al., 2006) using OrthoMCL v2.0.9 (Li et al., 2003). The phylogenetic inference and divergence time estimation were analyzed with conserved single-copy orthologs. Divergence times of *R. lucieae* and reference species were estimated by running two Markov chain Monte Carlo chains for 10,000,000 generations using BEAST2 v2.6.3 (Bouckaert et al., 2014) with the partitioned alignment and JTT amino-acid exchange rate matrix from IQ-TREE v2.2.0 (Chernomor et al., 2016) and ModelFinder (Kalyaanamoorthy et al., 2017), discarding the first 10 % of trees as post-burn-in. The divergence time estimation was calibrated using the oldest fossil records within Rosaceae (113.0–100.5 Mya; Peppe et al., 2008) and Eurosids (145.0–100.5 Mya; Ward and Jenney, 1899; Mellon et al., 1963). Based on identified orthologous groups and estimated divergence time, gene family evolution in *R. lucieae* and other rosaceous species was inferred using CAFE v5.1.0 (Mendes et al., 2020), identifying gene families with significant family size change (*p* < 0.01) as rapidly evolving gene families. To infer the significantly evolved gene functions in *R. lucieae*, GO and KO enrichment analyses on expanded or contracted gene families were performed using the R package clusterProfiler v4.4.4 (Wu et al., 2021).

The genomic synteny among the genome of *R. lucieae* and the reference genomes of its closely related species in Rosoideae (i.e., *R. chinensis*, *R. rugosa*, and *F. vesca*) was analyzed to infer the structural genomic variation within their evolution. The pairwise genetic homologies between species were assessed using blastp, and homology-based collinear blocks were identified using MCScanX (Wang et al., 2012). The collinearity blocks between species were visualized using SynVisio (Bandi and Gutwin, 2020). To detect the gene duplications within the genome of *R. lucieae*, the paralogous collinear blocks were also identified using blastp and MCscanX.

To understand the sequence divergences of homologous genes and their duplications within the Rosoideae lineage, Ks within paralogs of a species or orthologs between a pair of species were calculated using ParaAT v2.0 (Zhang et al., 2012), incorporating sequence alignments from MAFFT and Ks calculation from KaKs_Calculator v3.0 (Zhang, 2022). For Ks calculation, pairs of homologous genes were identified with percent identity scores over 70 % and percent positive scores over 80 % in all-to-all pairwise blastp comparisons of genes in each orthologous group. Particularly, the Ks of homologous pairs of genes annotated with the GO term for ‘response to salt stress’ or its child terms were calculated to understand the evolution of salt stress-related genes in *R. lucieae*.

### Gene duplication and divergence analysis

The duplications and divergence of genes in rapidly evolving gene families or annotated with gene functions involved in salt tolerance were explored to elucidate the influence of gene duplication and diversification in the evolution of salt tolerance of *R. lucieae*. The genes responsible for salt tolerance were compiled based on the broad investigation of plant salt-stress resistance (Botella et al., 2005; Munns, 2005; Gandhi and Oelmüller, 2023). The candidate genes were identified by their functional annotation based on the UniProtKB/Swiss-Prot and NCBI nr databases (E-value ≤ 1E−5), followed by the protein family and domain searches using HMMsearch v3.4 (Eddy, 2011) based on the Pfam database (E-value ≤ 1E−3). For example, ion transport protein family (PF00520) and cyclic nucleotide-binding domain (PF00027) were searched for the identification of CNGC1 genes, and E1-E2 ATPase (PF00122) or cation transport ATPase (P-type) (PF13246) families and cation transporting ATPase C-terminus (PF00689) or N-terminus (PF00690) domains were searched for the identification of ACA and ECA genes.

Among Rosoideae species, the genealogies of CNGC1 and ACA genes, of which the numbers of copies were notably increased in *R. lucieae*, were respectively inferred using IQ- TREE based on maximum-likelihood approaches with the JTT+F+R5 (CNGC1) and JTT+I+G4 (ACAs) substitution models selected by ModelFinder, after aligning amino-acid sequences using MAFFT. The robustness of each tree was assessed by 5000 replicates of bootstrap approximation using UFBoot2 (Hoang et al., 2018). The structural motifs enriched in Rosoideae CNGC1 or ACAs were identified using STREME v5.5.7 (Bailey, 2021), with the minimum motif width of 6 aa and the *p*-value cutoff of 0.01. The pairwise synteny of orthologous genes was identified using blastp by finding the best match between a pair of species.

## ACKNOWLEDGEMENTS

This study was supported by the National Research Foundation of Korea [grant number NRF-2017H1A2A1045787] to the first author.

## AUTHOR CONTRIBUTIONS

J.-H.J., Y.S. and S.-C.K. designed and coordinated the study. J.-H.J. and S.-C.K. collected the materials. J.-H.J., M.J. and Y.S. analyzed molecular data. J.-H.J. and S.-C.K. wrote the manuscript. The manuscript was reviewed and accepted by all the authors.

## DATA AVAILABILITY

Molecular sequencing data and genome assembly generated in this study were deposited into NCBI under BioProject accession PRJNA1069305.

## SUPPLEMENTARY DATA

Supplementary data is available at online in the Supplemental Material section.

**Fig. S1.** *K*-mer distribution in the short-read sequencing data of *Rosa lucieae*.

**Fig. S2.** Syntenic dotplot of two *R. lucieae* assemblies.

**Fig. S3.** Pairwise genome synteny of *R. lucieae* and its close relatives

**Fig. S4.** Comparison of pairwise Ks distribution of orthologous genes among the Rosoideae species.

**Fig. S5.** KEGG pathway enrichment analysis on *Rosa lucieae* genes within the rapidly evolving gene families based on the gene family evolution analysis.

**Fig. S6.** Duplication and divergence of CNGC1 genes within Rosoideae.

**Fig. S7.** Duplication of candidates of salt tolerance-related genes in *R. lucieae*.

**Table S1**. Summary of whole-genome and transcriptome sequencing of *R. lucieae* and subsequent read preprocessing.

**Table S2.** Statistics of the de novo genome assembly of *R. lucieae*.

**Table S3.** Statistics of the genome scaffolding of *R. lucieae* assembly.

**Table S4.** Repeat sequences within the genome of *R. lucieae*.

**Table S5.** Statistics of functional annotation of protein-coding genes from the genome of *R. lucieae*.

**Table S6.** List of the reference genomes and their gene models used for the homology-based gene prediction of *R. lucieae*.

**Table S7.** List of gene ontologies enriched in rapidly evolving gene families of *R. lucieae*.

**Table S8.** List of candidate genes related to salt tolerance in *R. lucieae*.

