## Supplemental Figures and Tables for "Genomic insights into the adaptive evolution of the memorial rose (*Rosa lucieae*; Rosaceae) in saline environments"

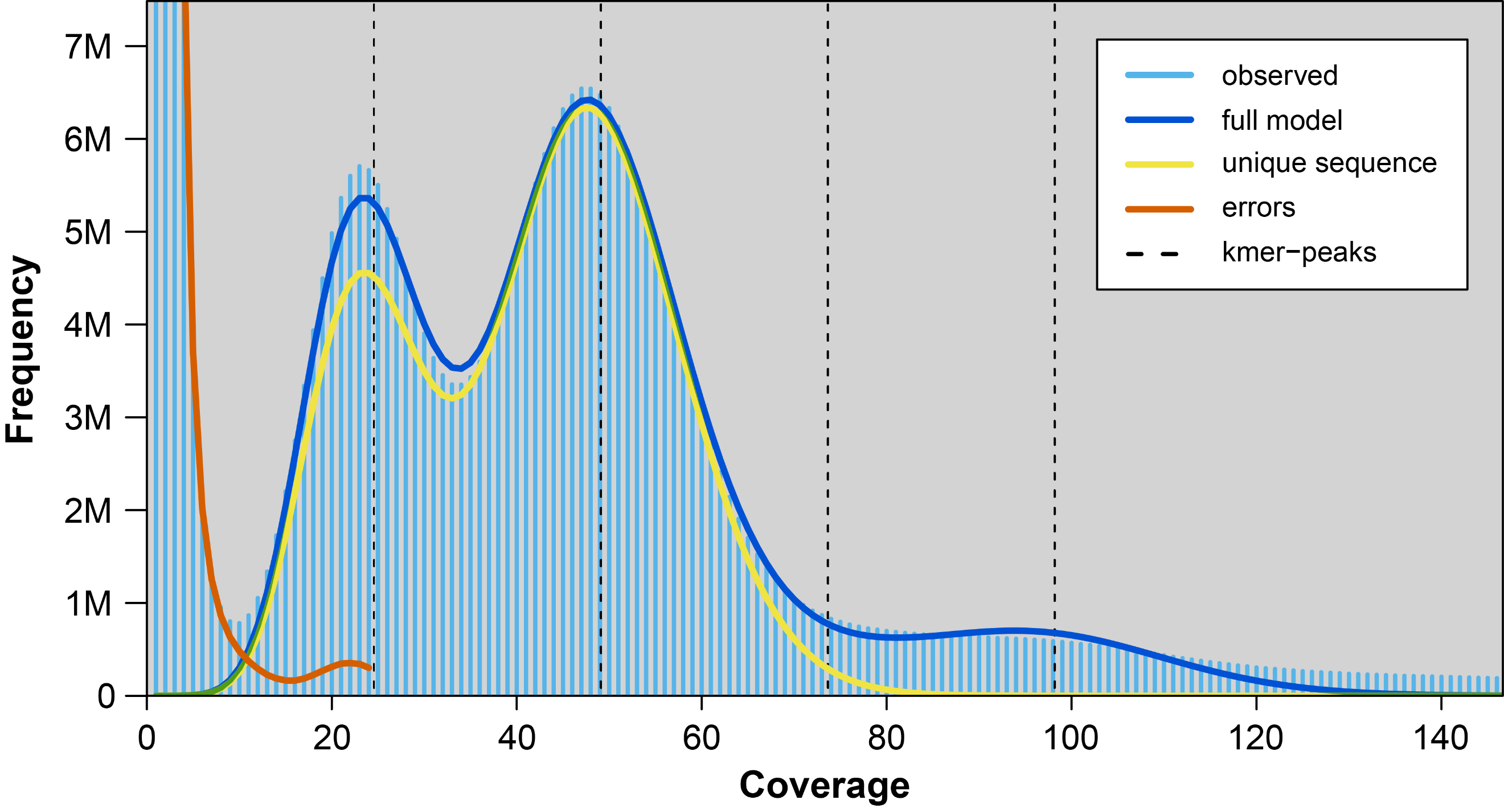

### Fig. S1. ***K*-mer distribution (*k* = 17) in the short-read sequencing data of *Rosa lucieae***.

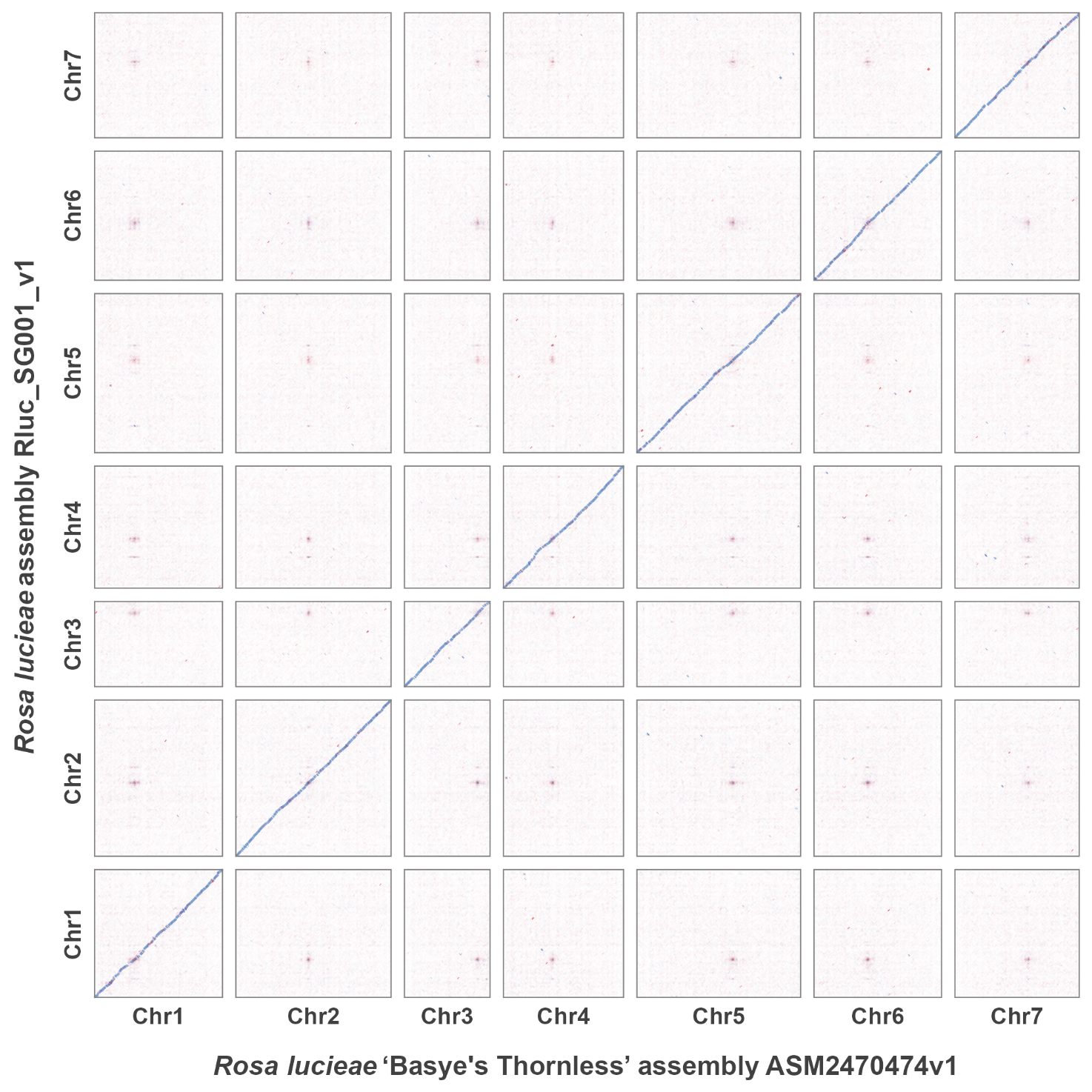

### Fig. S2. **Syntenic dotplot of two *Rosa lucieae* assemblies**, the wild *R. lucieae* assembly (Rluc_SG001_v1) from the present study and the *R. lucieae* ‘Basye's Thornless’ assembly (ASM2470474v1). Colored blocks indicate the collinear blocks, where the blue ones represent the blocks with parallel strands, and the red ones represents the blocks with antiparallel strands.

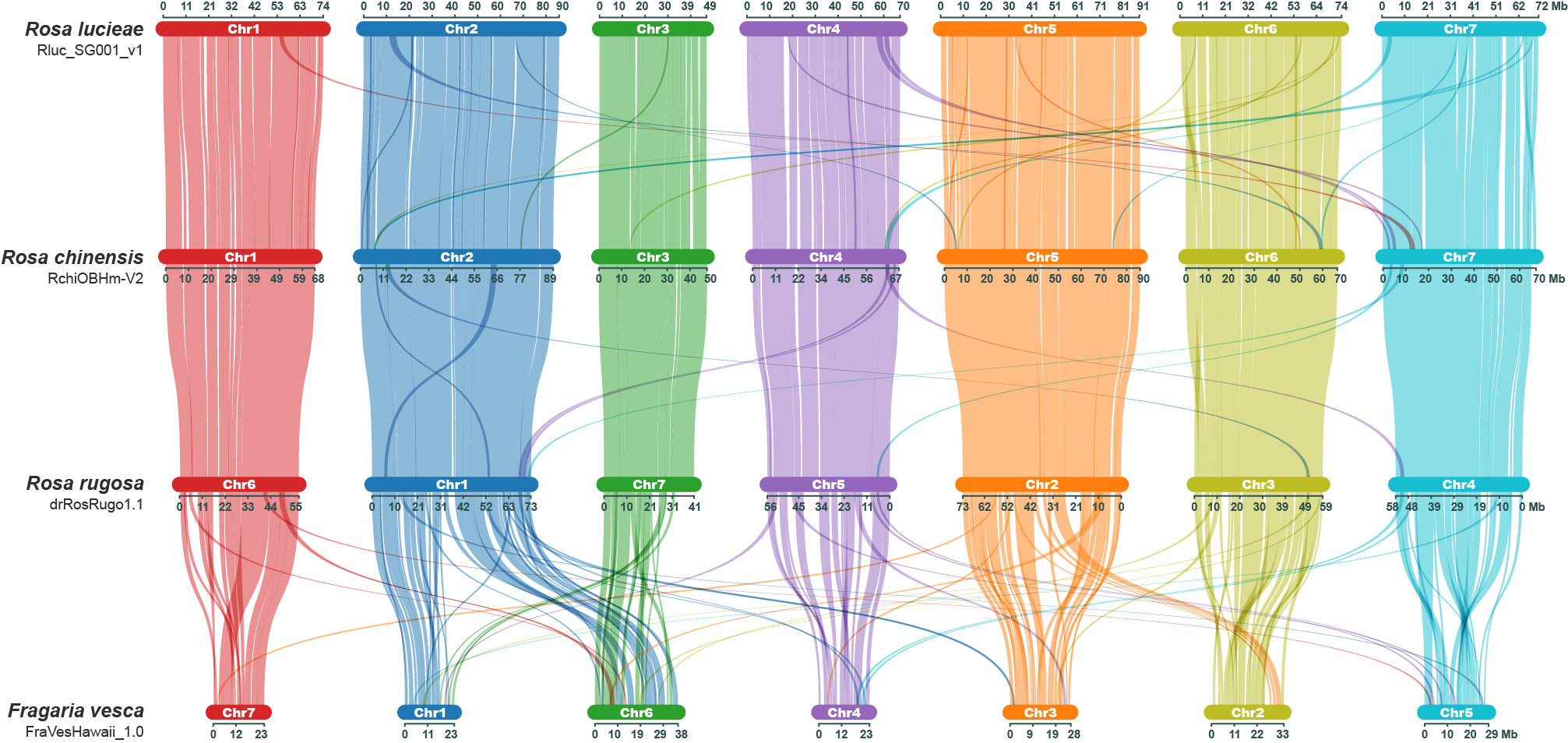

### Fig. S3. **Pairwise genome synteny of *Rosa lucieae* and its close relatives (*Rosa chinensis*, *R. rugosa* and *Fragaria vesca*)**. Each colored link represents the relationship of collinear blocks.

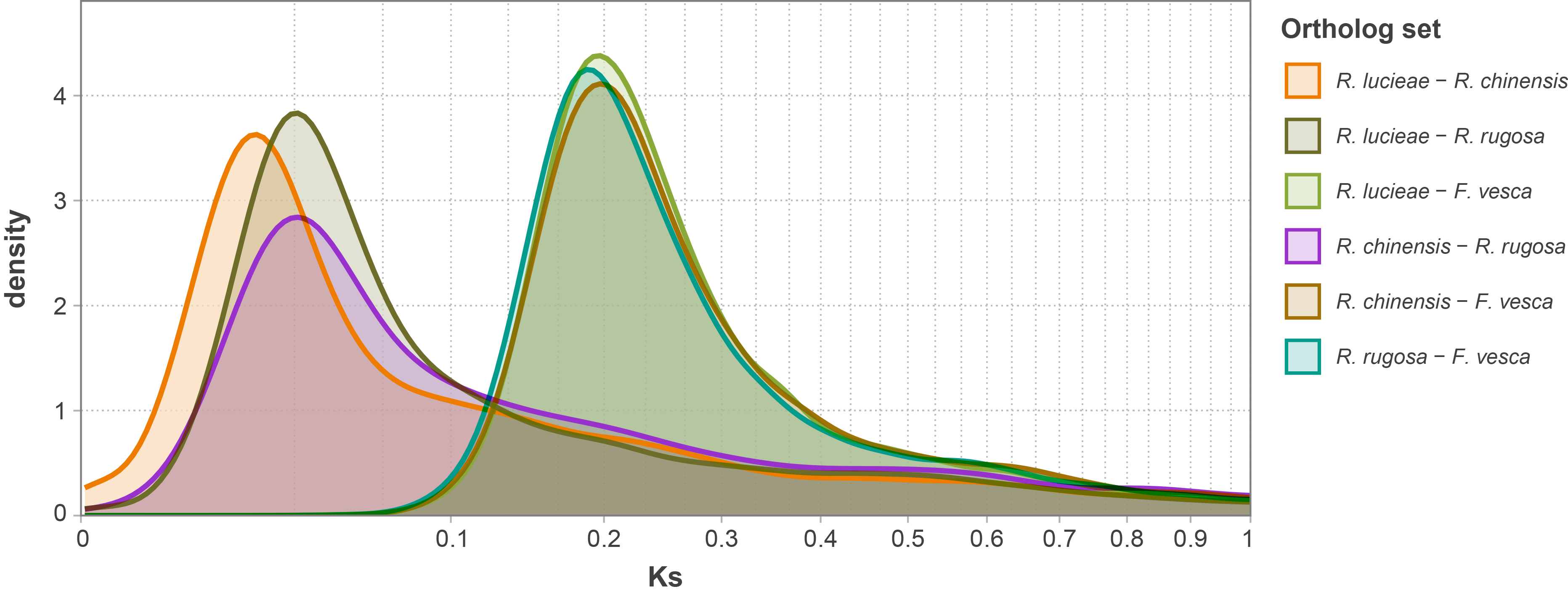

### Fig. S4. Comparison of pairwise Ks distribution of orthologous genes among the Rosoideae species.

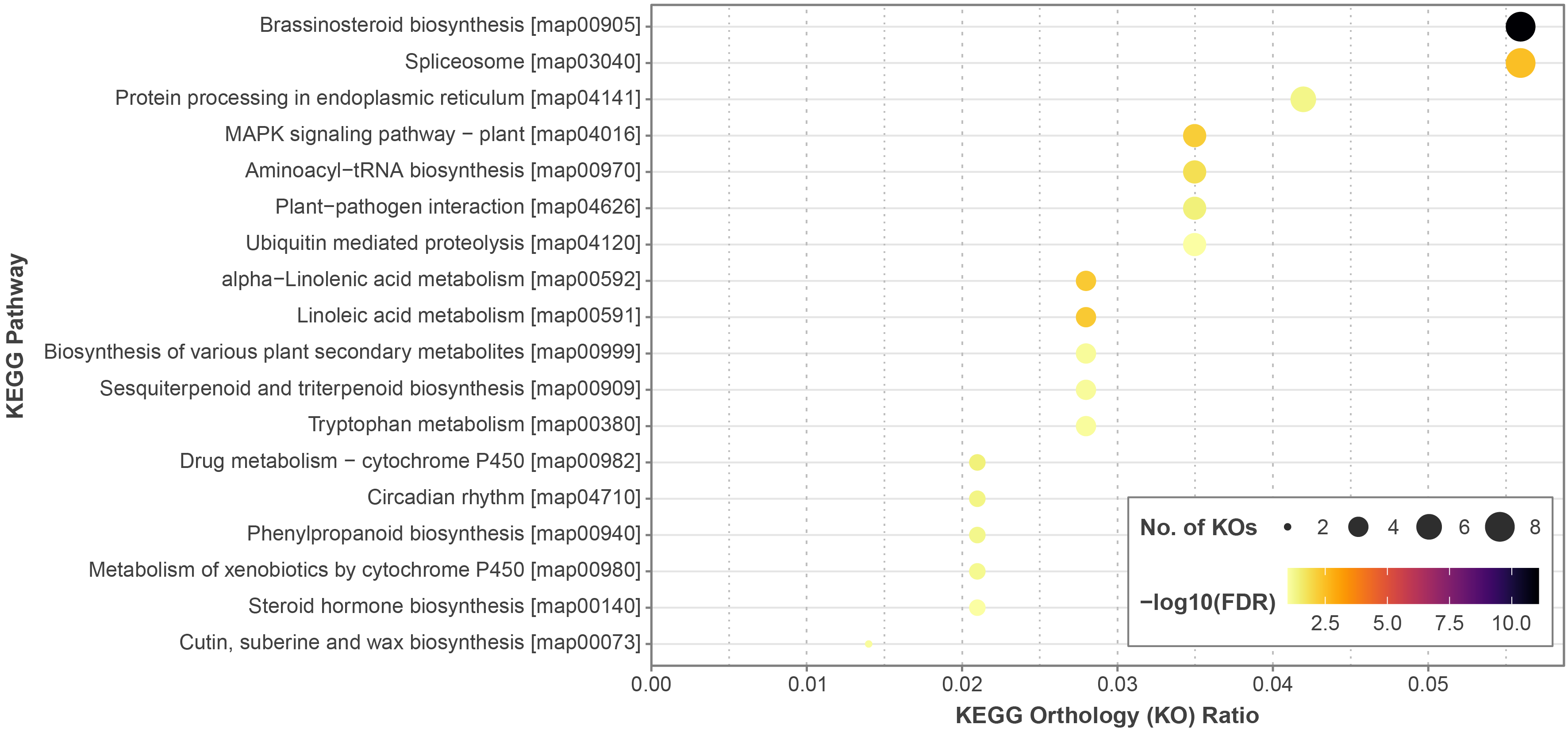

### Fig. S5. KEGG pathway enrichment analysis on KEGG orthologies of *Rosa lucieae* genes within the rapidly evolving gene families based on the gene family evolution analysis. KO: KEGG orthology; FDR: false discovery rate.

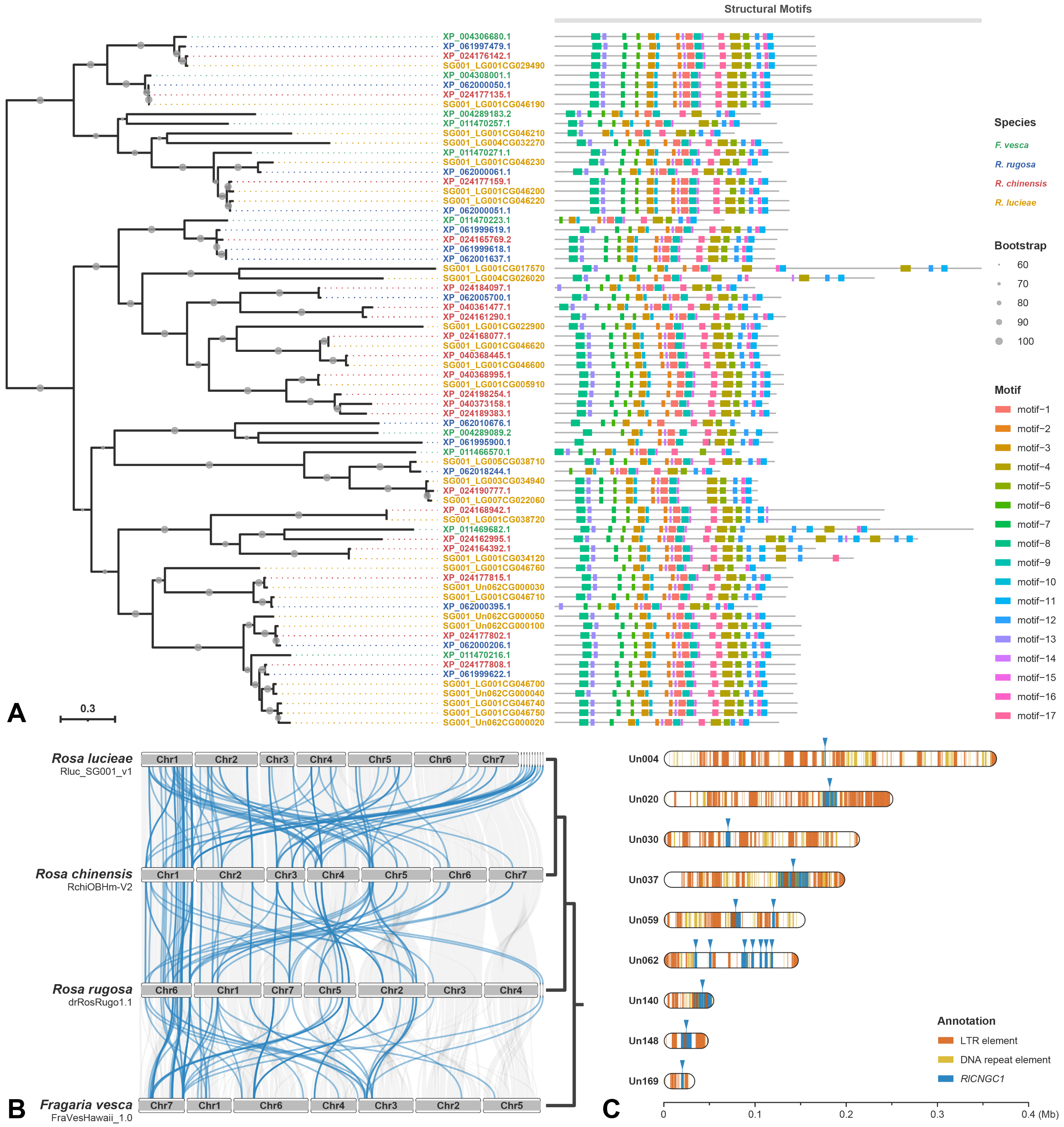

### Fig. S6. **Duplication and divergence of CNGC1 genes within Rosoideae.** A) **Unrooted gene tree (left) and the structural motifs (right) of CNGC1 genes which are significantly annotated with ion transport protein family (PF00520) and cyclic nucleotide-binding domain (PF00027).** B) **Ideogram of the four Rosoideae species showing synteny of CNGC1 genes between species. Blue lines represent the synteny relationships of orthologous CNGC1 gene pairs, and gray lines represent the synteny relationships of genomic collinear blocks. The cladogram on the right side represents the species relationships.** C) **Distribution map of long terminal repeats (LTRs), DNA repeat elements and CNGC1 genes within unplaced contigs of genome assembly of *Rosa lucieae* in this study**.

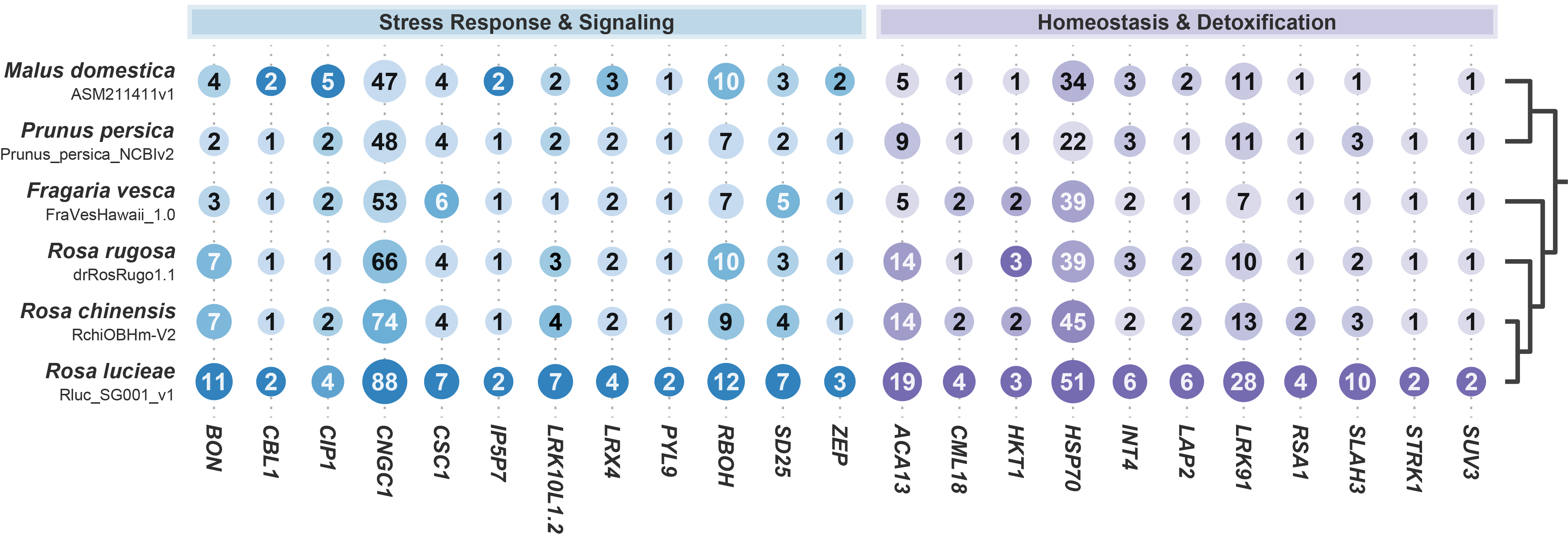

### Fig. S7. **Duplication of candidates of salt tolerance-related genes in *Rosa lucieae***. The bubble plot shows the copy numbers of candidate genes within the genomes of six rosaceous species, where the x-axis represents the candidate genes and the y-axis represents the genome assembly of each rosaceous species. The numbers in bubbles and sizes of bubbles represent the gene copy numbers, and the color shades of bubbles represent the normalized gene copy numbers in each gene. The blue plot includes the genes related to salt stress response and signaling, and the purple plot includes the genes related to homeostasis and detoxification. The cladogram on the right side represents the species relationships.

### Table S1. Summary of whole-genome and transcriptome sequencing of *Rosa lucieae* and subsequent read preprocessing.

| **Sequencing type** | **Sequencing platform** | **SRA accession** | **Description** | **No. of reads** | **Total read length (bp)** |
| --- | --- | --- | --- | --- | --- |
| Whole-genome sequencing | Illumina HiSeq  (Paired-end) | SRR27756703 | Raw sequence data | 168,651,380 | 25,466,358,380 |
|  |  |  | Quality-trimmed sequence reads | 167,947,240 | 24,974,320,045 |
|  |  |  | Contaminant-removed sequence reads | 154,712,198 | 23,024,594,713 |
|  |  | SRR27756704 | Raw sequence data | 205,978,888 | 31,102,812,088 |
|  |  |  | Quality-trimmed sequence reads | 203,449,492 | 30,002,717,168 |
|  |  |  | Contaminant-removed sequence reads | 187,934,712 | 27,734,408,577 |
|  | PacBio Sequel | SRR27756705 | Raw sequence data | 2,686,861 | 28,678,671,898 |
|  |  |  | Quality-trimmed sequence subreads | 3,523,501 | 28,641,998,351 |
| Transcriptome sequencing  (young leaves) | Illumina NovaSeq  (Paired-end) | SRR28392801 | Raw sequence data | 89,038,100 | 8,992,848,100 |
|  |  |  | Quality-trimmed sequence reads | 88,800,612 | 8,939,306,603 |
| Transcriptome sequencing  (flower buds) | Illumina NovaSeq  (Paired-end) | SRR28392802 | Raw sequence data | 88,807,358 | 8,969,543,158 |
|  |  |  | Quality-trimmed sequence reads | 88,484,096 | 8,902,264,907 |

### Table S2. Statistics of the *de novo* genome assembly of *Rosa lucieae*.

| **Feature** | **Primary assembly** | **Curated assembly** | **Draft assembly** |
| --- | --- | --- | --- |
| Number of contigs | 2067 | 1356 | 1356 |
| Total assembly size (bp) | 619,473,022 | 544,857,524 | 544,538,961 |
| Maximum contig length (bp) | 5,598,383 | 5,598,383 | 5,597,648 |
| Minimum contig length (bp) | 20,132 | 20,554 | 20,495 |
| Average contig length (bp) | 299,697 | 401,812 | 401,577 |
| Contig N50 (bp) | 546,059 | 636,224 | 635,886 |
| GC content (%) | 38.71 | 38.71 | 38.71 |

### Table S3. Statistics of the genome scaffolding of *Rosa lucieae* assembly.

| **Assembly** | **Total assembly  size (bp)** | **No. of  anchored contigs** | **Average length of  anchored contigs (bp)** | **N50 of  anchored contigs (bp)** |
| --- | --- | --- | --- | --- |
| Chr1 | 73,525,455 | 114 | 644,861.01 | 922,537 |
| Chr2 | 89,780,063 | 251 | 357,589.89 | 568,416 |
| Chr3 | 49,192,837 | 102 | 482,183.70 | 678,222 |
| Chr4 | 70,308,602 | 146 | 481,466.45 | 627,178 |
| Chr5 | 91,372,125 | 211 | 432,943.72 | 665,206 |
| Chr6 | 74,308,781 | 174 | 426,962.53 | 614,409 |
| Chr7 | 71,928,429 | 171 | 420,534.67 | 639,071 |
| **Assembly** | **Total assembly  size (bp)** | **No. of  unplaced contigs** | **Average length of  unplaced contigs (bp)** | **N50 of  unplaced contigs (bp)** |
| Unplaced contigs | 24,238,869 | 187 | 129,619.62 | 180,204 |

### Table S4. Repeat sequences within the genome of *Rosa lucieae*.

| **Classification** | **No. of elements** | **Repeat length (bp)** | **%age of sequences** |
| --- | --- | --- | --- |
| Interspersed repeats | 507,559 | 263,440,226 | 48.37 |
| SINEs | 4956 | 592,247 | 0.11 |
| ALUs | 0 | 0 | 0.00 |
| MIRs | 0 | 0 | 0.00 |
| LINEs | 27,650 | 14,834,227 | 2.72 |
| LINE1 | 26,053 | 14,176,236 | 2.60 |
| LINE2 | 649 | 187,053 | 0.03 |
| L3/CR1 | 0 | 0 | 0.00 |
| LTR elements | 120,190 | 142,520,358 | 26.17 |
| ERVL | 298 | 58,743 | 0.01 |
| ERVL-MaLRs | 0 | 0 | 0.00 |
| ERV_classI | 171 | 82,750 | 0.02 |
| ERV_classII | 0 | 0 | 0.00 |
| DNA elements | 127,356 | 40,873,481 | 7.50 |
| hAT-Charlie | 0 | 0 | 0.00 |
| TcMar-Tigger | 0 | 0 | 0.00 |
| Unclassified | 227,407 | 64,619,913 | 11.86 |
| Small RNA | 5230 | 685,123 | 0.13 |
| Satellites | 1401 | 872,509 | 0.16 |
| Simple repeats | 142,409 | 5,793,536 | 1.06 |
| Low complexity | 25,050 | 1,259,795 | 0.23 |

### Table S5. Statistics of functional annotation of protein-coding genes within the genome of *Rosa lucieae*.

|  | **nr** | **GO** | **KEGG** | **EggNOG** | **Pfam** | **Any  database** |
| --- | --- | --- | --- | --- | --- | --- |
| No. of  annotated genes | 36,870 | 33,795 | 30,378 | 35,439 | 33,012 | 36,873 |
| Percentage  (%) | 99.98 | 91.64 | 82.38 | 96.10 | 89.52 | 99.99 |

### Table S6. List of the reference genomes and their gene models used for the homology-based gene prediction of *Rosa lucieae*.

| **Species (Taxonomic classification)** | |  | **RefSeq accession ID** | **No. of  protein-coding genes** | **No. of proteins** |
| --- | --- | --- | --- | --- | --- |
| *Rosa chinensis* | (Rosoideae; Rosaceae; Rosales; Rosids) |  | GCF_002994745.2 | 30,924 | 48,188 |
| *Rosa rugosa* | (Rosoideae; Rosaceae; Rosales; Rosids) |  | GCF_958449725.1 | 29,146 | 42,829 |
| *Fragaria vesca* | (Rosoideae; Rosaceae; Rosales; Rosids) |  | GCF_000184155.1 | 22,383 | 31,387 |
| *Prunus persica* | (Amygdaloideae; Rosaceae; Rosales; Rosids) |  | GCF_000346465.2 | 23,135 | 32,595 |
| *Prunus avium* | (Amygdaloideae; Rosaceae; Rosales; Rosids) |  | GCF_002207925.1 | 25,841 | 35,009 |
| *Prunus dulcis* | (Amygdaloideae; Rosaceae; Rosales; Rosids) |  | GCF_902201215.1 | 23,151 | 33,326 |
| *Prunus mume* | (Amygdaloideae; Rosaceae; Rosales; Rosids) |  | GCF_000346735.1 | 23,423 | 29,705 |
| *Malus domestica* | (Amygdaloideae; Rosaceae; Rosales; Rosids) |  | GCF_002114115.1 | 35,926 | 52,036 |
| *Pyrus bretschneideri* | (Amygdaloideae; Rosaceae; Rosales; Rosids) |  | GCF_000315295.1 | 34,996 | 47,086 |
| *Quercus lobata* | (Fagaceae; Fagales; Rosids) |  | GCF_001633185.2 | 36,703 | 53,226 |
| *Glycine max* | (Fabaceae; Fabales; Rosids) |  | GCF_000004515.6 | 47,065 | 74,248 |
| *Glycine soja* | (Fabaceae; Fabales; Rosids) |  | GCF_004193775.1 | 47,201 | 69,277 |
| *Populus trichocarpa* | (Salicaceae; Malpighiales; Rosids) |  | GCF_000002775.4 | 31,641 | 51,717 |
| *Arabidopsis thaliana* | (Brassicaceae; Brassicales; Rosids) |  | GCF_000001735.4 | 27,562 | 48,265 |
| *Vitis vinifera* | (Vitaceae; Vitales; Rosids) |  | GCF_000003745.3 | 25,834 | 41,208 |

### Table S7. List of representative gene ontologies (GOs) significantly enriched within the rapidly evolving gene families of *Rosa lucieae*.

| **Category** | **GO ID** | **GO term** | **Gene ratio** | ***p*-value** | | **Adjusted *p*** | | **FDR** |
| --- | --- | --- | --- | --- | --- | --- | --- | --- |
| **Salinity-related GOs** | |  |  |  | |  | |  |
| BP | GO:0071474 | cellular hyperosmotic response | 11/1889 | 9.62E−12 | | 2.74E−10 | | 1.91E−10 |
| BP | GO:0071475 | cellular hyperosmotic salinity response | 10/1889 | 1.10E−10 | | 2.69E−09 | | 1.88E−09 |
| BP | GO:0071462 | cellular response to water stimulus | 16/1889 | 1.33E−06 | | 0.000015 | | 0.000010 |
| BP | GO:0042631 | cellular response to water deprivation | 19/1889 | 1.66E−06 | | 0.000018 | | 0.000013 |
| BP | GO:0042538 | hyperosmotic salinity response | 19/1889 | 0.000815 | | 0.004158 | | 0.002904 |
| BP | GO:0071470 | cellular response to osmotic stress | 14/1889 | 0.001356 | | 0.006597 | | 0.004608 |
| BP | GO:0006972 | hyperosmotic response | 19/1889 | 0.002412 | | 0.011036 | | 0.007708 |
| BP | GO:0009415 | response to water | 53/1889 | 0.005888 | | 0.023822 | | 0.016638 |
| BP | GO:0071472 | cellular response to salt stress | 10/1889 | 0.007590 | | 0.029913 | | 0.020892 |
| **Homeostasis-related GOs** | |  |  |  | |  | |  |
| BP | GO:0030104 | obsolete water homeostasis | 32/1889 | 9.87E−18 | | 5.02E−16 | | 3.51E−16 |
| BP | GO:0055075 | potassium ion homeostasis | 27/1889 | 2.46E−13 | | 8.42E−12 | | 5.88E−12 |
| BP | GO:0042044 | fluid transport | 26/1889 | 2.18E−10 | | 4.81E−09 | | 3.36E−09 |
| BP | GO:0006833 | water transport | 26/1889 | 9.89E−10 | | 2.08E−08 | | 1.45E−08 |
| BP | GO:0055065 | obsolete metal ion homeostasis | 38/1889 | 3.87E−07 | | 4.61E−06 | | 3.22E−06 |
| BP | GO:0055067 | obsolete monovalent inorganic cation homeostasis | 27/1889 | 4.38E−07 | | 5.15E−06 | | 3.59E−06 |
| BP | GO:0050801 | monoatomic ion homeostasis | 54/1889 | 8.67E−07 | | 9.71E−06 | | 6.78E−06 |
| BP | GO:0090332 | stomatal closure | 12/1889 | 0.000035 | | 0.000279 | | 0.000195 |
| BP | GO:0055080 | monoatomic cation homeostasis | 39/1889 | 0.000073 | | 0.000505 | | 0.000353 |
| BP | GO:0098771 | inorganic ion homeostasis | 39/1889 | 0.000420 | | 0.002352 | | 0.001643 |
| BP | GO:0050891 | multicellular organismal-level water homeostasis | 6/1889 | 0.001048 | | 0.005192 | | 0.003626 |
| BP | GO:0050878 | regulation of body fluid levels | 6/1889 | 0.008125 | | 0.031874 | | 0.022261 |
| BP | GO:0048871 | multicellular organismal-level homeostasis | 6/1889 | 0.015075 | | 0.053284 | | 0.037214 |
| **Ca^2+^-related GOs** | |  |  |  | |  | |  |
| BP | GO:0070588 | calcium ion transmembrane transport | 46/1889 | 2.16E−17 | | 1.07E−15 | | 7.46E−16 |
| BP | GO:0006816 | calcium ion transport | 46/1889 | 1.87E−16 | | 8.50E−15 | | 5.93E−15 |
| MF | GO:0015085 | calcium ion transmembrane transporter activity | 46/1910 | 1.92E−18 | | 4.20E−17 | | 2.20E−17 |
| MF | GO:0005388 | P-type calcium transporter activity | 33/1910 | 3.22E−18 | | 6.80E−17 | | 3.57E−17 |
| MF | GO:0005544 | calcium-dependent phospholipid binding | 18/1910 | 4.79E−09 | | 3.86E−08 | | 2.02E−08 |
| MF | GO:0005227 | calcium-activated cation channel activity | 7/1910 | 0.000146 | | 0.000515 | | 0.000270 |
| MF | GO:0005262 | calcium channel activity | 15/1910 | 0.002503 | | 0.006808 | | 0.003572 |
| **Signaling-related GOs** | |  |  |  |  | |  | |
| BP | GO:0009695 | jasmonic acid biosynthetic process | 40/1889 | 1.31E−23 | | 2.19E−21 | | 1.53E−21 |
| BP | GO:0042445 | hormone metabolic process | 91/1889 | 4.61E−21 | | 4.30E−19 | | 3.00E−19 |
| BP | GO:0010268 | brassinosteroid homeostasis | 27/1889 | 2.93E−12 | | 8.94E−11 | | 6.25E−11 |
| BP | GO:0016128 | phytosteroid metabolic process | 27/1889 | 9.02E−12 | | 2.61E−10 | | 1.82E−10 |
| BP | GO:0009696 | salicylic acid metabolic process | 22/1889 | 1.79E−11 | | 4.85E−10 | | 3.39E−10 |
| BP | GO:0016131 | brassinosteroid metabolic process | 27/1889 | 1.84E−11 | | 4.90E−10 | | 3.42E−10 |
| BP | GO:1901653 | cellular response to peptide | 20/1889 | 4.97E−11 | | 1.24E−09 | | 8.69E−10 |
| BP | GO:0009687 | abscisic acid metabolic process | 18/1889 | 9.57E−10 | | 2.03E−08 | | 1.42E−08 |
| BP | GO:0071731 | response to nitric oxide | 18/1889 | 2.38E−09 | | 4.55E−08 | | 3.18E−08 |
| BP | GO:0009741 | response to brassinosteroid | 41/1889 | 7.99E−09 | | 1.40E−07 | | 9.76E−08 |
| BP | GO:0046345 | abscisic acid catabolic process | 12/1889 | 1.02E−08 | | 1.75E−07 | | 1.22E−07 |
| BP | GO:0042446 | hormone biosynthetic process | 48/1889 | 2.61E−08 | | 4.07E−07 | | 2.84E−07 |
| BP | GO:0070301 | cellular response to hydrogen peroxide | 13/1889 | 3.77E−08 | | 5.65E−07 | | 3.95E−07 |
| BP | GO:0009926 | auxin polar transport | 25/1889 | 9.70E−08 | | 1.32E−06 | | 9.25E−07 |
| BP | GO:0042542 | response to hydrogen peroxide | 33/1889 | 4.69E−07 | | 5.43E−06 | | 3.79E−06 |
| BP | GO:0009694 | jasmonic acid metabolic process | 21/1889 | 2.07E−06 | | 0.000022 | | 0.000015 |
| BP | GO:0007166 | cell surface receptor signaling pathway | 65/1889 | 0.000017 | | 0.000149 | | 0.000104 |
| BP | GO:0071215 | cellular response to abscisic acid stimulus | 32/1889 | 0.000055 | | 0.000392 | | 0.000274 |
| BP | GO:0010928 | regulation of auxin mediated signaling pathway | 18/1889 | 0.000117 | | 0.000789 | | 0.000551 |
| BP | GO:0071396 | cellular response to lipid | 50/1889 | 0.000162 | | 0.001025 | | 0.000716 |
| BP | GO:0043200 | response to amino acid | 17/1889 | 0.000732 | | 0.003769 | | 0.002632 |
| BP | GO:0009692 | ethylene metabolic process | 7/1889 | 0.000900 | | 0.004540 | | 0.003171 |
| BP | GO:0009686 | gibberellin biosynthetic process | 18/1889 | 0.001495 | | 0.007173 | | 0.005010 |

BP: biological process; MF: molecular function; CC: cellular component; FDR: false discovery rate.

**Table S7.** *Continued*.

| **Category** | **GO ID** | **GO term** | **Gene ratio** | ***p*-value** | **Adjusted *p*** | **FDR** |
| --- | --- | --- | --- | --- | --- | --- |
| **Signaling-related GOs** *(continued)* | | |  |  |  |  |
| BP | GO:0071375 | cellular response to peptide hormone stimulus | 8/1889 | 0.001573 | 0.007483 | 0.005226 |
| BP | GO:0060918 | auxin transport | 17/1889 | 0.002432 | 0.011094 | 0.007748 |
| BP | GO:0009850 | auxin metabolic process | 12/1889 | 0.002973 | 0.013385 | 0.009348 |
| BP | GO:0009693 | ethylene biosynthetic process | 8/1889 | 0.003107 | 0.013874 | 0.009690 |
| BP | GO:0071370 | cellular response to gibberellin stimulus | 18/1889 | 0.003144 | 0.013990 | 0.009771 |
| BP | GO:1901652 | response to peptide | 8/1889 | 0.005608 | 0.023021 | 0.016078 |
| BP | GO:0034614 | cellular response to reactive oxygen species | 13/1889 | 0.005649 | 0.023132 | 0.016156 |
| BP | GO:0006694 | steroid biosynthetic process | 23/1889 | 0.005690 | 0.023246 | 0.016235 |
| BP | GO:0009685 | gibberellin metabolic process | 18/1889 | 0.006643 | 0.026618 | 0.018590 |
| BP | GO:0009914 | hormone transport | 15/1889 | 0.011270 | 0.041955 | 0.029302 |
| BP | GO:0009753 | response to jasmonic acid | 44/1889 | 0.011683 | 0.043171 | 0.030151 |
| BP | GO:0009688 | abscisic acid biosynthetic process | 6/1889 | 0.012420 | 0.045731 | 0.031939 |
| MF | GO:0001653 | peptide receptor activity | 28/1910 | 1.05E−15 | 1.68E−14 | 8.83E−15 |
| MF | GO:0042277 | peptide binding | 34/1910 | 2.52E−08 | 1.79E−07 | 9.41E−08 |
| MF | GO:0017046 | peptide hormone binding | 11/1910 | 8.93E−06 | 0.000038 | 0.000020 |
| MF | GO:0038023 | signaling receptor activity | 55/1910 | 0.005323 | 0.013083 | 0.006863 |
| MF | GO:0005102 | signaling receptor binding | 24/1910 | 0.022821 | 0.049482 | 0.025958 |
| MF | GO:0051428 | peptide hormone receptor binding | 2/1910 | 0.041849 | 0.083973 | 0.044052 |
| **Metabolite-related GOs** | |  |  |  |  |  |
| BP | GO:0046274 | lignin catabolic process | 67/1889 | 9.35E−69 | 1.57E−65 | 1.10E−65 |
| BP | GO:0018874 | benzoate metabolic process | 18/1889 | 1.77E−20 | 1.29E−18 | 9.0E−19 |
| BP | GO:1901615 | organic hydroxy compound metabolic process | 87/1889 | 1.76E−18 | 1.13E−16 | 7.92E−17 |
| BP | GO:0046482 | para-aminobenzoic acid metabolic process | 18/1889 | 2.87E−18 | 1.56E−16 | 1.09E−16 |
| BP | GO:0016134 | saponin metabolic process | 20/1889 | 1.85E−16 | 8.50E−15 | 5.93E−15 |
| BP | GO:0018958 | phenol-containing compound metabolic process | 29/1889 | 5.21E−14 | 1.99E−12 | 1.39E−12 |
| BP | GO:0009820 | alkaloid metabolic process | 32/1889 | 7.87E−14 | 2.94E−12 | 2.05E−12 |
| BP | GO:0016107 | sesquiterpenoid catabolic process | 12/1889 | 5.39E−12 | 1.59E−10 | 1.11E−10 |
| BP | GO:0043290 | apocarotenoid catabolic process | 12/1889 | 5.39E−12 | 1.59E−10 | 1.11E−10 |
| BP | GO:0016128 | phytosteroid metabolic process | 27/1889 | 9.02E−12 | 2.61E−10 | 1.82E−10 |
| BP | GO:0043288 | apocarotenoid metabolic process | 18/1889 | 2.13E−10 | 4.77E−09 | 3.33E−09 |
| BP | GO:1902644 | tertiary alcohol metabolic process | 18/1889 | 2.13E−10 | 4.77E−09 | 3.33E−09 |
| BP | GO:0006066 | alcohol metabolic process | 35/1889 | 6.73E−09 | 1.20E−07 | 8.39E−08 |
| BP | GO:0009808 | lignin metabolic process | 22/1889 | 1.85E−08 | 3.05E−07 | 2.13E−07 |
| BP | GO:0009809 | lignin biosynthetic process | 32/1889 | 1.97E−08 | 3.21E−07 | 2.24E−07 |
| BP | GO:0006714 | sesquiterpenoid metabolic process | 18/1889 | 2.62E−08 | 4.07E−07 | 2.84E−07 |
| BP | GO:0010023 | proanthocyanidin biosynthetic process | 12/1889 | 7.47E−08 | 1.04E−06 | 7.24E−07 |
| BP | GO:0046246 | terpene biosynthetic process | 19/1889 | 8.58E−08 | 1.18E−06 | 8.25E−07 |
| BP | GO:0016144 | S-glycoside biosynthetic process | 15/1889 | 1.37E−07 | 1.83E−06 | 1.28E−06 |
| BP | GO:0019758 | glycosinolate biosynthetic process | 15/1889 | 1.37E−07 | 1.83E−06 | 1.28E−06 |
| BP | GO:0042537 | benzene-containing compound metabolic process | 18/1889 | 1.42E−07 | 1.86E−06 | 1.30E−06 |
| BP | GO:0009072 | aromatic amino acid metabolic process | 25/1889 | 4.20E−07 | 4.96E−06 | 3.47E−06 |
| BP | GO:1901657 | glycosyl compound metabolic process | 39/1889 | 4.50E−07 | 5.24E−06 | 3.66E−06 |
| BP | GO:0019761 | glucosinolate biosynthetic process | 15/1889 | 6.09E−07 | 6.95E−06 | 4.86E−06 |
| BP | GO:0006721 | terpenoid metabolic process | 47/1889 | 2.39E−06 | 0.000025 | 0.000018 |
| BP | GO:0016135 | saponin biosynthetic process | 8/1889 | 3.28E−06 | 0.000033 | 0.000023 |
| BP | GO:0016114 | terpenoid biosynthetic process | 40/1889 | 5.72E−06 | 0.000055 | 0.000039 |
| BP | GO:0006720 | isoprenoid metabolic process | 50/1889 | 0.000014 | 0.000125 | 0.000087 |
| BP | GO:1901659 | glycosyl compound biosynthetic process | 23/1889 | 0.000015 | 0.000133 | 0.000093 |
| BP | GO:0046189 | phenol-containing compound biosynthetic process | 11/1889 | 0.000136 | 0.000888 | 0.000620 |
| BP | GO:0009698 | phenylpropanoid metabolic process | 29/1889 | 0.000420 | 0.002352 | 0.001643 |
| BP | GO:1901617 | organic hydroxy compound biosynthetic process | 32/1889 | 0.000429 | 0.002394 | 0.001672 |
| BP | GO:0016143 | S-glycoside metabolic process | 16/1889 | 0.000623 | 0.003268 | 0.002283 |
| BP | GO:0019757 | glycosinolate metabolic process | 16/1889 | 0.000623 | 0.003268 | 0.002283 |
| BP | GO:0006631 | fatty acid metabolic process | 42/1889 | 0.000664 | 0.003450 | 0.002409 |
| BP | GO:0008299 | isoprenoid biosynthetic process | 36/1889 | 0.000809 | 0.004140 | 0.002891 |
| BP | GO:0044550 | secondary metabolite biosynthetic process | 46/1889 | 0.000921 | 0.004631 | 0.003234 |
| BP | GO:0019748 | secondary metabolic process | 56/1889 | 0.002205 | 0.010116 | 0.007065 |

BP: biological process; MF: molecular function; CC: cellular component; FDR: false discovery rate.

**Table S7.** *Continued*.

| **Category** | **GO ID** | **GO term** | **Gene ratio** | ***p*-value** | **Adjusted *p*** | **FDR** |
| --- | --- | --- | --- | --- | --- | --- |
| **Metabolite-related GOs** *(continued)* | | |  |  |  |  |
| BP | GO:0043289 | apocarotenoid biosynthetic process | 6/1889 | 0.003840 | 0.016488 | 0.011516 |
| BP | GO:1902645 | tertiary alcohol biosynthetic process | 6/1889 | 0.003840 | 0.016488 | 0.011516 |
| BP | GO:0072330 | monocarboxylic acid biosynthetic process | 35/1889 | 0.004242 | 0.018079 | 0.012626 |
| BP | GO:0019760 | glucosinolate metabolic process | 16/1889 | 0.005914 | 0.023867 | 0.016669 |
| BP | GO:0006722 | triterpenoid metabolic process | 11/1889 | 0.007582 | 0.029913 | 0.020892 |
| BP | GO:0016101 | diterpenoid metabolic process | 18/1889 | 0.019834 | 0.068240 | 0.047659 |
| MF | GO:0046527 | glucosyltransferase activity | 74/1910 | 5.48E−20 | 1.77E−18 | 9.26E−19 |
| MF | GO:0035251 | UDP-glucosyltransferase activity | 70/1910 | 2.74E−19 | 6.71E−18 | 3.52E−18 |
| MF | GO:0008194 | UDP-glycosyltransferase activity | 74/1910 | 1.29E−12 | 1.62E−11 | 8.48E−12 |
| **Protein repair-related GOs** | | |  |  |  |  |
| BP | GO:0042026 | protein refolding | 24/1889 | 1.00E−10 | 2.48E−09 | 1.73E−09 |
| BP | GO:0051085 | chaperone cofactor-dependent protein refolding | 25/1889 | 1.13E−08 | 1.92E−07 | 1.34E−07 |
| BP | GO:0030091 | protein repair | 8/1889 | 0.000940 | 0.004698 | 0.003281 |
| MF | GO:0140662 | ATP-dependent protein folding chaperone | 51/1910 | 1.19E−15 | 1.87E−14 | 9.80E−15 |
| MF | GO:0031072 | heat shock protein binding | 24/1910 | 0.000039 | 0.000155 | 0.000081 |
| MF | GO:0031625 | ubiquitin protein ligase binding | 42/1910 | 0.012457 | 0.028769 | 0.015092 |
| **Root- & Growth-related GOs** | | |  |  |  |  |
| BP | GO:0090708 | specification of plant organ axis polarity | 60/1889 | 2.29E−46 | 1.28E−43 | 8.94E−44 |
| BP | GO:1903224 | regulation of endodermal cell differentiation | 26/1889 | 6.84E−23 | 8.83E−21 | 6.17E−21 |
| BP | GO:0048507 | meristem development | 93/1889 | 1.02E−22 | 1.14E−20 | 7.97E−21 |
| BP | GO:0007492 | endoderm development | 26/1889 | 9.97E−21 | 7.61E−19 | 5.31E−19 |
| BP | GO:0010073 | meristem maintenance | 76/1889 | 2.20E−18 | 1.37E−16 | 9.55E−17 |
| BP | GO:2000280 | regulation of root development | 67/1889 | 6.50E−18 | 3.41E−16 | 2.38E−16 |
| BP | GO:0009630 | gravitropism | 20/1889 | 0.000264 | 0.001591 | 0.001111 |
| BP | GO:0010071 | root meristem specification | 8/1889 | 0.000277 | 0.001646 | 0.001149 |
| BP | GO:0010274 | hydrotropism | 8/1889 | 0.000387 | 0.002195 | 0.001533 |
| BP | GO:0009934 | regulation of meristem structural organization | 9/1889 | 0.001740 | 0.008207 | 0.005732 |
| BP | GO:0010102 | lateral root morphogenesis | 17/1889 | 0.003708 | 0.016253 | 0.011352 |
| BP | GO:0048830 | adventitious root development | 5/1889 | 0.009036 | 0.034482 | 0.024083 |
| BP | GO:0010075 | regulation of meristem growth | 9/1889 | 0.014839 | 0.052674 | 0.036788 |
| CC | GO:0048226 | Casparian strip | 64/637 | 3.97E−57 | 1.01E−54 | 6.74E−55 |
| **Defense-related GOs** | |  |  |  |  |  |
| BP | GO:0002238 | response to molecule of fungal origin | 23/1889 | 1.16E−07 | 1.57E−06 | 1.09E−06 |
| BP | GO:0080027 | response to herbivore | 17/1889 | 1.67E−07 | 2.15E−06 | 1.50E−06 |
| BP | GO:0002239 | response to oomycetes | 29/1889 | 2.53E−07 | 3.14E−06 | 2.19E−06 |
| BP | GO:0002229 | defense response to oomycetes | 28/1889 | 1.66E−06 | 0.000018 | 0.000013 |
| BP | GO:0009626 | plant-type hypersensitive response | 63/1889 | 2.18E−06 | 0.000023 | 0.000016 |
| BP | GO:0031349 | positive regulation of defense response | 32/1889 | 0.000016 | 0.000143 | 0.000100 |
| BP | GO:0042545 | cell wall modification | 28/1889 | 0.000021 | 0.000176 | 0.000123 |
| BP | GO:0052545 | callose localization | 15/1889 | 0.000024 | 0.000195 | 0.000136 |
| BP | GO:0052386 | cell wall thickening | 13/1889 | 0.000058 | 0.000409 | 0.000286 |
| BP | GO:0050776 | regulation of immune response | 40/1889 | 0.000121 | 0.000816 | 0.000570 |
| BP | GO:0052543 | callose deposition in cell wall | 14/1889 | 0.000125 | 0.000839 | 0.000586 |
| BP | GO:0052542 | defense response by callose deposition | 11/1889 | 0.000136 | 0.000888 | 0.000620 |
| BP | GO:0002684 | positive regulation of immune system process | 24/1889 | 0.000250 | 0.001517 | 0.001059 |
| BP | GO:0034050 | symbiont-induced defense-related programmed cell death | 20/1889 | 0.000301 | 0.001780 | 0.001243 |
| BP | GO:0052482 | defense response by cell wall thickening | 9/1889 | 0.000537 | 0.002896 | 0.002022 |
| BP | GO:0045089 | positive regulation of innate immune response | 20/1889 | 0.000628 | 0.003287 | 0.002296 |
| BP | GO:0002213 | defense response to insect | 12/1889 | 0.001903 | 0.008924 | 0.006232 |
| BP | GO:1900367 | positive regulation of defense response to insect | 6/1889 | 0.002113 | 0.009749 | 0.006809 |
| BP | GO:0009624 | response to nematode | 25/1889 | 0.002955 | 0.013337 | 0.009314 |
| BP | GO:0052544 | defense response by callose deposition in cell wall | 9/1889 | 0.003150 | 0.013990 | 0.009771 |
| BP | GO:0050778 | positive regulation of immune response | 20/1889 | 0.003599 | 0.015860 | 0.011077 |
| BP | GO:0002758 | innate immune response-activating signaling pathway | 11/1889 | 0.005089 | 0.021150 | 0.014771 |
| BP | GO:0045088 | regulation of innate immune response | 27/1889 | 0.009541 | 0.036242 | 0.025312 |
| BP | GO:0002218 | activation of innate immune response | 11/1889 | 0.018901 | 0.065299 | 0.045606 |

BP: biological process; MF: molecular function; CC: cellular component; FDR: false discovery rate.

### Table S8. List of candidate genes related to salt tolerance in *Rosa lucieae*.

| **Gene ID** | **Protein**  **length** | **NCBI nr blastp** |  | **UniProtKB/Swiss-Prot blastp** |  |
| --- | --- | --- | --- | --- | --- |
|  |  | **Gene description** | **E-value** | **Gene description** | **E-value** |
| ***RlACA13*** |  |  |  |  |  |
| SG001_LG002CG052780 | 1075 | putative calcium-transporting ATPase 13, plasma membrane-type | 0 | Putative calcium-transporting ATPase 13, plasma membrane-type | 1.68E−141 |
| SG001_LG002CG052800 | 1024 | putative calcium-transporting ATPase 13, plasma membrane-type | 0 | Calcium-transporting ATPase 12, plasma membrane-type | 1.67E−128 |
| SG001_LG004CG000370 | 1033 | putative calcium-transporting ATPase 13, plasma membrane-type | 0 | Putative calcium-transporting ATPase 13, plasma membrane-type | 0 |
| SG001_LG004CG001750 | 987 | putative calcium-transporting ATPase 13, plasma membrane-type | 0 | Putative calcium-transporting ATPase 13, plasma membrane-type | 0 |
| SG001_LG004CG001770 | 1003 | putative calcium-transporting ATPase 13, plasma membrane-type | 0 | Calcium-transporting ATPase 12, plasma membrane-type | 0 |
| SG001_LG004CG001790 | 966 | putative calcium-transporting ATPase 13, plasma membrane-type | 0 | Putative calcium-transporting ATPase 13, plasma membrane-type | 0 |
| SG001_LG004CG001800 | 1040 | putative calcium-transporting ATPase 13, plasma membrane-type | 0 | Putative calcium-transporting ATPase 13, plasma membrane-type | 0 |
| SG001_LG004CG001810 | 1040 | putative calcium-transporting ATPase 13, plasma membrane-type | 0 | Putative calcium-transporting ATPase 13, plasma membrane-type | 0 |
| SG001_LG004CG001830 | 1035 | putative calcium-transporting ATPase 13, plasma membrane-type | 0 | Putative calcium-transporting ATPase 13, plasma membrane-type | 0 |
| SG001_LG004CG001860 | 881 | putative calcium-transporting ATPase 13, plasma membrane-type | 0 | Putative calcium-transporting ATPase 13, plasma membrane-type | 0 |
| SG001_LG004CG001870 | 1011 | putative calcium-transporting ATPase 13, plasma membrane-type | 0 | Putative calcium-transporting ATPase 13, plasma membrane-type | 0 |
| SG001_LG004CG001890 | 327 | putative calcium-transporting ATPase 13, plasma membrane-type | 3.73E−177 | Putative calcium-transporting ATPase 13, plasma membrane-type | 1.06E−78 |
| SG001_LG004CG001940 | 1020 | putative calcium-transporting ATPase 13, plasma membrane-type | 0 | Putative calcium-transporting ATPase 13, plasma membrane-type | 0 |
| SG001_LG004CG001960 | 1036 | putative calcium-transporting ATPase 13, plasma membrane-type | 0 | Putative calcium-transporting ATPase 13, plasma membrane-type | 0 |
| SG001_LG004CG001980 | 1033 | putative calcium-transporting ATPase 13, plasma membrane-type | 0 | Putative calcium-transporting ATPase 13, plasma membrane-type | 0 |
| SG001_LG005CG000800 | 1031 | putative calcium-transporting ATPase 13, plasma membrane-type | 0 | Putative calcium-transporting ATPase 13, plasma membrane-type | 1.87E−176 |
| SG001_LG005CG045750 | 962 | putative calcium-transporting ATPase 13, plasma membrane-type | 0 | Putative calcium-transporting ATPase 13, plasma membrane-type | 0 |
| SG001_LG005CG045760 | 1017 | putative calcium-transporting ATPase 13, plasma membrane-type | 0 | Putative calcium-transporting ATPase 13, plasma membrane-type | 0 |
| SG001_LG005CG045770 | 985 | putative calcium-transporting ATPase 13, plasma membrane-type | 0 | Putative calcium-transporting ATPase 13, plasma membrane-type | 0 |
| ***RlBON1*** |  |  |  |  |  |
| SG001_LG002CG045690 | 574 | protein BONZAI 1-like | 0 | Protein BONZAI 1 | 0 |
| SG001_LG007CG009120 | 574 | protein BONZAI 1-like | 0 | Protein BONZAI 1 | 0 |
| SG001_LG007CG009160 | 574 | protein BONZAI 1-like | 0 | Protein BONZAI 1 | 0 |
| ***RlBON3*** |  |  |  |  |  |
| SG001_LG002CG033130 | 585 | protein BONZAI 3 isoform X1 | 0 | Protein BONZAI 3 | 0 |
| SG001_LG003CG004470 | 583 | protein BONZAI 3-like isoform X1 | 0 | Protein BONZAI 3 | 0 |
| SG001_LG003CG004480 | 583 | protein BONZAI 3 isoform X1 | 0 | Protein BONZAI 3 | 0 |
| SG001_LG003CG004490 | 589 | putative C2 domain, von Willebrand factor, type A, copine, protein BONZAI | 0 | Protein BONZAI 3 | 0 |
| SG001_LG003CG004500 | 590 | protein BONZAI 3 isoform X1 | 0 | Protein BONZAI 3 | 0 |
| SG001_LG003CG004510 | 590 | protein BONZAI 3 isoform X1 | 0 | Protein BONZAI 3 | 0 |
| SG001_LG003CG004520 | 602 | protein BONZAI 3 isoform X1 | 0 | Protein BONZAI 3 | 0 |
| SG001_LG004CG017010 | 587 | protein BONZAI 3 isoform X1 | 0 | Protein BONZAI 3 | 0 |
| ***RlCBL1*** |  |  |  |  |  |
| SG001_LG004CG009540 | 213 | calcineurin B-like protein 1 | 2.27E−152 | Calcineurin B-like protein 1 | 3.30E−130 |
| SG001_LG007CG021450 | 213 | calcineurin B-like protein 1 | 2.27E−152 | Calcineurin B-like protein 1 | 3.30E−130 |

**Table S8.** *Continued*.

| **Gene ID** | **Protein**  **length** | **NCBI nr blastp** |  | **UniProtKB/Swiss-Prot blastp** |  |
| --- | --- | --- | --- | --- | --- |
|  |  | **Gene description** | **E-value** | **Gene description** | **E-value** |
| ***RlCIP1*** |  |  |  |  |  |
| SG001_LG004CG007910 | 932 | COP1-interactive protein 1-like | 0 | COP1-interactive protein 1 | 1.35E−33 |
| SG001_LG004CG007920 | 1686 | COP1-interactive protein 1 | 0 | COP1-interactive protein 1 | 1.24E−124 |
| SG001_LG007CG014340 | 1680 | COP1-interactive protein 1 | 0 | COP1-interactive protein 1 | 1.06E−126 |
| SG001_LG007CG014350 | 932 | COP1-interactive protein 1-like | 0 | COP1-interactive protein 1 | 2.15E−33 |
| ***RlCML18*** |  |  |  |  |  |
| SG001_LG001CG047460 | 171 | probable calcium-binding protein CML18 | 9.73E−119 | Probable calcium-binding protein CML18 | 4.15E−71 |
| SG001_LG005CG044600 | 163 | probable calcium-binding protein CML18 | 9.77E−115 | Probable calcium-binding protein CML17 | 9.65E−87 |
| SG001_LG007CG024810 | 163 | probable calcium-binding protein CML18 | 9.77E−115 | Probable calcium-binding protein CML17 | 9.65E−87 |
| SG001_LG007CG024850 | 163 | probable calcium-binding protein CML18 | 9.77E−115 | Probable calcium-binding protein CML17 | 9.65E−87 |
| ***RlCNGC1*** |  |  |  |  |  |
| SG001_LG001CG005910 | 634 | cyclic nucleotide-gated ion channel 1-like isoform X1 | 0 | Cyclic nucleotide-gated ion channel 1 | 3.03E−135 |
| SG001_LG001CG005930 | 942 | cyclic nucleotide-gated ion channel 1-like | 0 | Cyclic nucleotide-gated ion channel 1 | 3.30E−84 |
| SG001_LG001CG008230 | 618 | cyclic nucleotide-gated ion channel 1-like | 0 | Cyclic nucleotide-gated ion channel 1 | 8.34E−93 |
| SG001_LG001CG013050 | 646 | cyclic nucleotide-gated ion channel 1-like | 0 | Cyclic nucleotide-gated ion channel 1 | 1.04E−87 |
| SG001_LG001CG014310 | 673 | cyclic nucleotide-gated ion channel 1-like | 0 | Cyclic nucleotide-gated ion channel 1 | 4.65E−130 |
| SG001_LG001CG017570 | 1180 | cyclic nucleotide-gated ion channel 1-like | 0 | Cyclic nucleotide-gated ion channel 1 | 5.06E−93 |
| SG001_LG001CG022900 | 589 | cyclic nucleotide-gated ion channel 1-like | 0 | Cyclic nucleotide-gated ion channel 1 | 4.96E−115 |
| SG001_LG001CG024380 | 620 | cyclic nucleotide-gated ion channel 1-like | 0 | Cyclic nucleotide-gated ion channel 1 | 2.43E−66 |
| SG001_LG001CG024430 | 623 | cyclic nucleotide-gated ion channel 1-like | 0 | Cyclic nucleotide-gated ion channel 1 | 3.09E−71 |
| SG001_LG001CG024460 | 262 | cyclic nucleotide-gated ion channel 1-like | 4.99E−179 | Cyclic nucleotide-gated ion channel 1 | 1.21E−39 |
| SG001_LG001CG024500 | 619 | cyclic nucleotide-gated ion channel 1-like | 0 | Cyclic nucleotide-gated ion channel 1 | 1.04E−77 |
| SG001_LG001CG024530 | 628 | cyclic nucleotide-gated ion channel 1-like | 0 | Cyclic nucleotide-gated ion channel 1 | 2.23E−76 |
| SG001_LG001CG027190 | 650 | cyclic nucleotide-gated ion channel 1-like | 0 | Cyclic nucleotide-gated ion channel 1 | 9.16E−87 |
| SG001_LG001CG029490 | 725 | cyclic nucleotide-gated ion channel 1-like | 0 | Cyclic nucleotide-gated ion channel 1 | 0 |
| SG001_LG001CG032280 | 724 | cyclic nucleotide-gated ion channel 1 | 0 | Cyclic nucleotide-gated ion channel 1 | 3.25E−89 |
| SG001_LG001CG032290 | 626 | cyclic nucleotide-gated ion channel 1 | 0 | Cyclic nucleotide-gated ion channel 1 | 5.38E−86 |
| SG001_LG001CG033420 | 605 | cyclic nucleotide-gated ion channel 1-like | 0 | Cyclic nucleotide-gated ion channel 1 | 3.76E−112 |
| SG001_LG001CG034120 | 827 | cyclic nucleotide-gated ion channel 1 isoform X1 | 0 | Cyclic nucleotide-gated ion channel 1 | 2.45E−137 |
| SG001_LG001CG034130 | 488 | cyclic nucleotide-gated ion channel 1-like isoform X1 | 0 | Cyclic nucleotide-gated ion channel 1 | 1.62E−83 |
| SG001_LG001CG034170 | 974 | cyclic nucleotide-gated ion channel 1-like isoform X1 | 0 | Cyclic nucleotide-gated ion channel 1 | 2.20E−97 |
| SG001_LG001CG038720 | 900 | cyclic nucleotide-gated ion channel 1-like | 0 | Cyclic nucleotide-gated ion channel 1 | 6.14E−120 |
| SG001_LG001CG046190 | 714 | cyclic nucleotide-gated ion channel 1-like | 0 | Cyclic nucleotide-gated ion channel 1 | 0 |
| SG001_LG001CG046200 | 621 | cyclic nucleotide-gated ion channel 1-like | 0 | Cyclic nucleotide-gated ion channel 1 | 0 |
| SG001_LG001CG046210 | 498 | cyclic nucleotide-gated ion channel 1-like | 0 | Cyclic nucleotide-gated ion channel 1 | 3.40E−124 |
| SG001_LG001CG046220 | 649 | cyclic nucleotide-gated ion channel 1-like | 0 | Cyclic nucleotide-gated ion channel 1 | 0 |

**Table S8.** *Continued*.

| **Gene ID** | **Protein**  **length** | **NCBI nr blastp** |  | **UniProtKB/Swiss-Prot blastp** |  |
| --- | --- | --- | --- | --- | --- |
|  |  | **Gene description** | **E-value** | **Gene description** | **E-value** |
| ***RlCNGC1*** *(continued)* |  |  |  |  |  |
| SG001_LG001CG046230 | 603 | cyclic nucleotide-gated ion channel 1-like | 0 | Cyclic nucleotide-gated ion channel 1 | 2.92E−178 |
| SG001_LG001CG046250 | 616 | cyclic nucleotide-gated ion channel 1-like | 0 | Cyclic nucleotide-gated ion channel 1 | 0 |
| SG001_LG001CG046600 | 589 | cyclic nucleotide-gated ion channel 1-like | 0 | Cyclic nucleotide-gated ion channel 1 | 4.75E−136 |
| SG001_LG001CG046620 | 617 | cyclic nucleotide-gated ion channel 1-like | 0 | Cyclic nucleotide-gated ion channel 1 | 1.01E−130 |
| SG001_LG001CG046630 | 646 | cyclic nucleotide-gated ion channel 1-like | 0 | Cyclic nucleotide-gated ion channel 1 | 3.24E−124 |
| SG001_LG001CG046650 | 603 | cyclic nucleotide-gated ion channel 1-like | 0 | Cyclic nucleotide-gated ion channel 1 | 3.11E−120 |
| SG001_LG001CG046700 | 671 | cyclic nucleotide-gated ion channel 1-like | 0 | Cyclic nucleotide-gated ion channel 1 | 6.17E−139 |
| SG001_LG001CG046710 | 640 | cyclic nucleotide-gated ion channel 1-like | 0 | Cyclic nucleotide-gated ion channel 1 | 1.48E−144 |
| SG001_LG001CG046740 | 672 | cyclic nucleotide-gated ion channel 1-like | 0 | Cyclic nucleotide-gated ion channel 1 | 6.91E−140 |
| SG001_LG001CG046750 | 671 | cyclic nucleotide-gated ion channel 1-like | 0 | Cyclic nucleotide-gated ion channel 1 | 4.20E−143 |
| SG001_LG001CG046760 | 641 | cyclic nucleotide-gated ion channel 1-like | 0 | Cyclic nucleotide-gated ion channel 1 | 2.85E−143 |
| SG001_LG002CG007830 | 278 | cyclic nucleotide-gated ion channel 1-like | 0 | Cyclic nucleotide-gated ion channel 1 | 2.05E−29 |
| SG001_LG002CG017230 | 691 | cyclic nucleotide-gated ion channel 1 | 0 | Cyclic nucleotide-gated ion channel 1 | 2.86E−120 |
| SG001_LG002CG018380 | 482 | cyclic nucleotide-gated ion channel 1 | 0 | Cyclic nucleotide-gated ion channel 1 | 1.95E−129 |
| SG001_LG002CG020780 | 690 | cyclic nucleotide-gated ion channel 1 | 0 | Cyclic nucleotide-gated ion channel 1 | 6.15E−119 |
| SG001_LG002CG027970 | 595 | cyclic nucleotide-gated ion channel 1-like | 0 | Cyclic nucleotide-gated ion channel 1 | 5.14E−62 |
| SG001_LG002CG028020 | 349 | cyclic nucleotide-gated ion channel 1-like | 1.02E−154 | Cyclic nucleotide-gated ion channel 1 | 5.01E−38 |
| SG001_LG002CG052320 | 249 | cyclic nucleotide-gated ion channel 1-like | 2.69E−137 | Cyclic nucleotide-gated ion channel 1 | 8.53E−16 |
| SG001_LG002CG052360 | 245 | cyclic nucleotide-gated ion channel 1-like | 1.81E−139 | Cyclic nucleotide-gated ion channel 1 | 7.45E−15 |
| SG001_LG002CG052370 | 275 | cyclic nucleotide-gated ion channel 1-like | 3.63E−137 | Cyclic nucleotide-gated ion channel 1 | 1.75E−20 |
| SG001_LG002CG052790 | 789 | cyclic nucleotide-gated ion channel 1 isoform X1 | 0 | Cyclic nucleotide-gated ion channel 1 | 3.66E−55 |
| SG001_LG002CG052810 | 829 | cyclic nucleotide-gated ion channel 1 isoform X1 | 0 | Cyclic nucleotide-gated ion channel 1 | 2.45E−100 |
| SG001_LG003CG013940 | 287 | cyclic nucleotide-gated ion channel 1-like | 5.07E−155 | Cyclic nucleotide-gated ion channel 1 | 7.02E−39 |
| SG001_LG003CG014090 | 570 | cyclic nucleotide-gated ion channel 1-like | 0 | Cyclic nucleotide-gated ion channel 1 | 2.65E−94 |
| SG001_LG003CG014120 | 509 | cyclic nucleotide-gated ion channel 1-like | 0 | Cyclic nucleotide-gated ion channel 1 | 6.62E−87 |
| SG001_LG003CG017190 | 600 | cyclic nucleotide-gated ion channel 1-like | 0 | Cyclic nucleotide-gated ion channel 1 | 2.22E−120 |
| SG001_LG003CG034720 | 939 | cyclic nucleotide-gated ion channel 1-like | 0 | Cyclic nucleotide-gated ion channel 1 | 9.54E−81 |
| SG001_LG003CG034940 | 563 | cyclic nucleotide-gated ion channel 1-like | 0 | Cyclic nucleotide-gated ion channel 1 | 2.44E−96 |
| SG001_LG004CG011310 | 551 | cyclic nucleotide-gated ion channel 1-like | 0 | Cyclic nucleotide-gated ion channel 1 | 4.04E−25 |
| SG001_LG004CG011470 | 625 | cyclic nucleotide-gated ion channel 1-like | 0 | Cyclic nucleotide-gated ion channel 1 | 1.05E−58 |
| SG001_LG004CG011540 | 647 | cyclic nucleotide-gated ion channel 1-like | 0 | Cyclic nucleotide-gated ion channel 1 | 1.05E−85 |
| SG001_LG004CG016110 | 631 | cyclic nucleotide-gated ion channel 1 | 0 | Cyclic nucleotide-gated ion channel 1 | 3.26E−113 |
| SG001_LG004CG026020 | 885 | cyclic nucleotide-gated ion channel 1-like | 0 | Cyclic nucleotide-gated ion channel 1 | 2.05E−114 |
| SG001_LG004CG032270 | 631 | cyclic nucleotide-gated ion channel 1-like | 0 | Cyclic nucleotide-gated ion channel 1 | 7.85E−143 |
| SG001_LG005CG009920 | 950 | cyclic nucleotide-gated ion channel 1-like | 0 | Cyclic nucleotide-gated ion channel 1 | 3.69E−109 |

**Table S8.** *Continued*.

| **Gene ID** | **Protein**  **length** | **NCBI nr blastp** |  | **UniProtKB/Swiss-Prot blastp** |  |
| --- | --- | --- | --- | --- | --- |
|  |  | **Gene description** | **E-value** | **Gene description** | **E-value** |
| ***RlCNGC1*** *(continued)* |  |  |  |  |  |
| SG001_LG005CG009940 | 950 | cyclic nucleotide-gated ion channel 1-like | 0 | Cyclic nucleotide-gated ion channel 1 | 9.83E−110 |
| SG001_LG005CG014100 | 1632 | cyclic nucleotide-gated ion channel 1 | 0 | Cyclic nucleotide-gated ion channel 1 | 1.05E−16 |
| SG001_LG005CG033120 | 626 | cyclic nucleotide-gated ion channel 1-like | 0 | Cyclic nucleotide-gated ion channel 1 | 3.37E−101 |
| SG001_LG005CG038710 | 609 | cyclic nucleotide-gated ion channel 1-like | 0 | Cyclic nucleotide-gated ion channel 1 | 1.27E−97 |
| SG001_LG005CG043040 | 545 | cyclic nucleotide-gated ion channel 1-like | 5.13E−155 | Cyclic nucleotide-gated ion channel 1 | 1.53E−79 |
| SG001_LG005CG046870 | 604 | cyclic nucleotide-gated ion channel 1-like isoform X1 | 0 | Cyclic nucleotide-gated ion channel 1 | 2.37E−124 |
| SG001_LG005CG046890 | 604 | cyclic nucleotide-gated ion channel 1-like | 0 | Cyclic nucleotide-gated ion channel 1 | 2.85E−120 |
| SG001_LG005CG058650 | 647 | cyclic nucleotide-gated ion channel 1 | 0 | Cyclic nucleotide-gated ion channel 1 | 1.66E−103 |
| SG001_LG006CG048330 | 600 | cyclic nucleotide-gated ion channel 1-like | 0 | Probable cyclic nucleotide-gated ion channel 10 | 1.95E−87 |
| SG001_LG007CG014840 | 381 | cyclic nucleotide-gated ion channel 1-like | 0 | Cyclic nucleotide-gated ion channel 1 | 1.24E−41 |
| SG001_LG007CG022060 | 563 | cyclic nucleotide-gated ion channel 1-like | 0 | Cyclic nucleotide-gated ion channel 1 | 5.82E−99 |
| SG001_LG007CG031850 | 535 | cyclic nucleotide-gated ion channel 1-like | 0 | Cyclic nucleotide-gated ion channel 1 | 2.77E−98 |
| SG001_Un004CG000080 | 206 | cyclic nucleotide-gated ion channel 1-like | 1.76E−105 | Putative cyclic nucleotide-gated ion channel 13 | 6.66E−22 |
| SG001_Un020CG000010 | 584 | cyclic nucleotide-gated ion channel 1-like | 0 | Cyclic nucleotide-gated ion channel 1 | 8.25E−114 |
| SG001_Un030CG000070 | 623 | cyclic nucleotide-gated ion channel 1-like | 0 | Cyclic nucleotide-gated ion channel 1 | 4.60E−87 |
| SG001_Un037CG000010 | 640 | cyclic nucleotide-gated ion channel 1-like | 0 | Cyclic nucleotide-gated ion channel 1 | 7.04E−87 |
| SG001_Un059CG000040 | 646 | cyclic nucleotide-gated ion channel 1-like | 0 | Cyclic nucleotide-gated ion channel 1 | 7.83E−125 |
| SG001_Un059CG000050 | 570 | cyclic nucleotide-gated ion channel 1-like | 0 | Cyclic nucleotide-gated ion channel 1 | 6.92E−86 |
| SG001_Un062CG000020 | 621 | cyclic nucleotide-gated ion channel 1-like | 0 | Cyclic nucleotide-gated ion channel 1 | 8.84E−128 |
| SG001_Un062CG000030 | 645 | cyclic nucleotide-gated ion channel 1-like | 0 | Cyclic nucleotide-gated ion channel 1 | 7.06E−136 |
| SG001_Un062CG000040 | 660 | cyclic nucleotide-gated ion channel 1-like | 0 | Cyclic nucleotide-gated ion channel 1 | 7.37E−138 |
| SG001_Un062CG000050 | 666 | cyclic nucleotide-gated ion channel 1-like | 0 | Cyclic nucleotide-gated ion channel 1 | 9.25E−143 |
| SG001_Un062CG000060 | 648 | cyclic nucleotide-gated ion channel 1-like | 0 | Cyclic nucleotide-gated ion channel 1 | 7.96E−122 |
| SG001_Un062CG000100 | 683 | cyclic nucleotide-gated ion channel 1-like | 0 | Cyclic nucleotide-gated ion channel 1 | 7.67E−140 |
| SG001_Un062CG000110 | 655 | cyclic nucleotide-gated ion channel 1-like | 0 | Cyclic nucleotide-gated ion channel 1 | 2.59E−125 |
| SG001_Un140CG000010 | 923 | cyclic nucleotide-gated ion channel 1-like | 0 | Cyclic nucleotide-gated ion channel 1 | 5.02E−84 |
| SG001_Un148CG000010 | 564 | cyclic nucleotide-gated ion channel 1-like | 0 | Cyclic nucleotide-gated ion channel 1 | 1.21E−34 |
| SG001_Un169CG000010 | 622 | cyclic nucleotide-gated ion channel 1-like | 0 | Cyclic nucleotide-gated ion channel 1 | 2.09E−105 |
| ***RlCSC1*** |  |  |  |  |  |
| SG001_LG001CG009600 | 669 | CSC1-like protein At1g32090 | 0 | CSC1-like protein At1g32090 | 0 |
| SG001_LG002CG054190 | 767 | CSC1-like protein At3g21620 | 0 | Calcium permeable stress-gated cation channel 1 | 0 |
| SG001_LG005CG001820 | 777 | calcium permeable stress-gated cation channel 1-like | 0 | Calcium permeable stress-gated cation channel 1 | 0 |
| SG001_LG006CG012210 | 747 | CSC1-like protein At3g21620 isoform X1 | 0 | Calcium permeable stress-gated cation channel 1 | 0 |
| SG001_LG006CG031890 | 688 | CSC1-like protein At1g32090 | 0 | CSC1-like protein At1g32090 | 0 |
| SG001_LG006CG034280 | 763 | CSC1-like protein At4g02900 | 0 | CSC1-like protein At4g02900 | 0 |
| SG001_LG007CG025890 | 833 | CSC1-like protein At1g32090 | 0 | CSC1-like protein At1g32090 | 0 |

**Table S8.** *Continued*.

| **Gene ID** | **Protein**  **length** | **NCBI nr blastp** |  | **UniProtKB/Swiss-Prot blastp** |  |
| --- | --- | --- | --- | --- | --- |
|  |  | **Gene description** | **E-value** | **Gene description** | **E-value** |
| ***RlHKT1*** |  |  |  |  |  |
| SG001_LG004CG008450 | 538 | sodium transporter HKT1-like | 0 | Sodium transporter HKT1 | 2.07E−160 |
| SG001_LG004CG008470 | 558 | sodium transporter HKT1-like | 0 | Sodium transporter HKT1 | 2.86E−169 |
| SG001_LG005CG041240 | 535 | sodium transporter HKT1-like | 0 | Sodium transporter HKT1 | 4.88E−166 |
| ***RlHSP70*** |  |  |  |  |  |
| SG001_LG001CG015730 | 697 | stromal 70 kDa heat shock-related protein, chloroplastic | 0 | Stromal 70 kDa heat shock-related protein, chloroplastic | 0 |
| SG001_LG001CG018820 | 649 | heat shock cognate 70 kDa protein 2 | 0 | Heat shock cognate 70 kDa protein | 0 |
| SG001_LG001CG020210 | 620 | heat shock cognate 70 kDa protein 2 | 0 | Heat shock cognate 70 kDa protein | 0 |
| SG001_LG001CG030370 | 365 | heat shock cognate 70 kDa protein-like | 0 | Heat shock cognate 70 kDa protein | 0 |
| SG001_LG001CG034260 | 512 | heat shock cognate 70 kDa protein-like | 0 | Heat shock cognate 70 kDa protein | 0 |
| SG001_LG001CG034430 | 411 | heat shock cognate 70 kDa protein-like | 0 | Heat shock 70 kDa protein 4 | 0 |
| SG001_LG001CG035670 | 649 | heat shock cognate 70 kDa protein 2 | 0 | Heat shock cognate 70 kDa protein | 0 |
| SG001_LG001CG035680 | 651 | heat shock cognate 70 kDa protein 2 | 0 | Heat shock cognate 70 kDa protein | 0 |
| SG001_LG001CG035790 | 475 | heat shock cognate 70 kDa protein-like | 0 | Heat shock cognate 70 kDa protein | 0 |
| SG001_LG001CG035810 | 521 | heat shock cognate 70 kDa protein-like | 0 | Heat shock 70 kDa protein 4 | 0 |
| SG001_LG001CG035830 | 528 | heat shock cognate 70 kDa protein-like | 0 | Heat shock 70 kDa protein 4 | 0 |
| SG001_LG001CG035850 | 530 | heat shock cognate 70 kDa protein-like | 0 | Heat shock cognate 70 kDa protein | 0 |
| SG001_LG001CG035870 | 526 | heat shock cognate 70 kDa protein-like | 0 | Heat shock cognate 70 kDa protein | 0 |
| SG001_LG001CG035880 | 518 | heat shock cognate 70 kDa protein-like | 0 | Heat shock cognate 70 kDa protein | 0 |
| SG001_LG001CG035890 | 533 | heat shock cognate 70 kDa protein-like | 0 | Heat shock cognate 70 kDa protein | 0 |
| SG001_LG001CG035900 | 523 | heat shock cognate 70 kDa protein-like | 0 | Heat shock cognate 70 kDa protein | 0 |
| SG001_LG001CG035910 | 533 | heat shock cognate 70 kDa protein-like | 0 | Heat shock cognate 70 kDa protein | 0 |
| SG001_LG001CG035920 | 533 | heat shock cognate 70 kDa protein-like | 0 | Heat shock cognate 70 kDa protein | 0 |
| SG001_LG001CG035930 | 524 | heat shock cognate 70 kDa protein-like | 0 | Heat shock cognate 70 kDa protein | 0 |
| SG001_LG002CG000870 | 654 | heat shock 70 kDa protein | 0 | Heat shock 70 kDa protein | 0 |
| SG001_LG002CG020060 | 654 | heat shock 70 kDa protein | 0 | Heat shock 70 kDa protein | 0 |
| SG001_LG002CG034010 | 619 | heat shock cognate 70 kDa protein 2-like | 0 | Heat shock cognate 70 kDa protein | 0 |
| SG001_LG002CG038850 | 678 | heat shock 70 kDa protein, mitochondrial | 0 | Heat shock 70 kDa protein, mitochondrial | 0 |
| SG001_LG003CG006010 | 661 | luminal-binding protein 5-like | 0 | Luminal-binding protein 5 | 0 |
| SG001_LG003CG014300 | 647 | heat shock 70 kDa protein 1 | 0 | Heat shock 70 kDa protein 1 | 0 |
| SG001_LG003CG014710 | 512 | heat shock cognate 70 kDa protein-like | 0 | Heat shock cognate 70 kDa protein | 0 |
| SG001_LG003CG014720 | 518 | heat shock cognate 70 kDa protein-like | 0 | Heat shock cognate 70 kDa protein | 0 |
| SG001_LG004CG000390 | 497 | putative Heat shock protein 70 family | 0 | Heat shock cognate 70 kDa protein 2 | 0 |
| SG001_LG004CG010260 | 618 | heat shock cognate 70 kDa protein 2 | 0 | Heat shock cognate 70 kDa protein 2 | 0 |
| SG001_LG004CG010380 | 587 | putative Heat shock protein 70 family | 0 | Heat shock cognate 70 kDa protein | 0 |
| SG001_LG004CG010410 | 619 | putative Heat shock protein 70 family | 0 | Heat shock cognate 70 kDa protein 2 | 0 |
| SG001_LG004CG010720 | 618 | putative Heat shock protein 70 family | 0 | Heat shock cognate 70 kDa protein 2 | 0 |

**Table S8.** *Continued*.

| **Gene ID** | **Protein**  **length** | **NCBI nr blastp** |  | **UniProtKB/Swiss-Prot blastp** |  |
| --- | --- | --- | --- | --- | --- |
|  |  | **Gene description** | **E-value** | **Gene description** | **E-value** |
| ***RlHSP70*** *(continued)* |  |  |  |  |  |
| SG001_LG004CG010760 | 618 | putative Heat shock protein 70 family | 0 | Heat shock cognate 70 kDa protein | 0 |
| SG001_LG004CG023310 | 573 | putative Heat shock protein 70 family | 0 | Heat shock cognate 70 kDa protein | 0 |
| SG001_LG005CG015140 | 624 | heat shock cognate 70 kDa protein 2 | 0 | Heat shock cognate 70 kDa protein | 0 |
| SG001_LG005CG015150 | 621 | putative Heat shock protein 70 family | 0 | Heat shock cognate 70 kDa protein | 0 |
| SG001_LG005CG055300 | 696 | stromal 70 kDa heat shock-related protein, chloroplastic | 0 | Stromal 70 kDa heat shock-related protein, chloroplastic | 0 |
| SG001_LG006CG001420 | 544 | putative Heat shock protein 70 family | 0 | Heat shock cognate 70 kDa protein | 0 |
| SG001_LG006CG001580 | 530 | heat shock cognate 70 kDa protein-like | 0 | Heat shock cognate 70 kDa protein | 0 |
| SG001_LG006CG002040 | 555 | heat shock cognate 70 kDa protein 2 | 0 | Heat shock cognate 70 kDa protein 2 | 0 |
| SG001_LG006CG002120 | 636 | heat shock cognate 70 kDa protein 2 | 0 | Heat shock cognate 70 kDa protein | 0 |
| SG001_LG006CG002150 | 562 | heat shock cognate 70 kDa protein 2 | 0 | Heat shock cognate 70 kDa protein 2 | 0 |
| SG001_LG006CG002190 | 550 | heat shock cognate 70 kDa protein 2 | 0 | Heat shock cognate 70 kDa protein | 0 |
| SG001_LG006CG002280 | 560 | heat shock cognate 70 kDa protein 2 | 0 | Heat shock cognate 70 kDa protein | 0 |
| SG001_LG006CG002300 | 529 | putative Heat shock protein 70 family | 0 | Heat shock cognate 70 kDa protein | 0 |
| SG001_LG006CG017590 | 676 | heat shock 70 kDa protein, mitochondrial | 0 | Heat shock 70 kDa protein, mitochondrial | 0 |
| SG001_LG007CG007260 | 666 | luminal-binding protein 5 | 0 | Luminal-binding protein 5 | 0 |
| SG001_LG007CG033370 | 696 | stromal 70 kDa heat shock-related protein, chloroplastic | 0 | Stromal 70 kDa heat shock-related protein, chloroplastic | 0 |
| SG001_LG007CG042500 | 651 | heat shock cognate 70 kDa protein 2 | 0 | Heat shock cognate 70 kDa protein | 0 |
| SG001_LG007CG044800 | 522 | heat shock cognate 70 kDa protein-like | 0 | Heat shock cognate 70 kDa protein | 0 |
| SG001_LG007CG044810 | 523 | heat shock cognate 70 kDa protein-like | 0 | Heat shock cognate 70 kDa protein | 0 |
| ***RlINT4*** |  |  |  |  |  |
| SG001_LG001CG038430 | 496 | inositol transporter 4-like | 0 | Inositol transporter 4 | 0 |
| SG001_LG001CG038460 | 367 | inositol transporter 4-like | 0 | Inositol transporter 4 | 1.71E−150 |
| SG001_LG004CG026480 | 480 | inositol transporter 4-like | 0 | Inositol transporter 4 | 0 |
| SG001_LG005CG000890 | 579 | inositol transporter 4-like | 0 | Inositol transporter 4 | 0 |
| SG001_LG005CG000900 | 581 | inositol transporter 4-like | 0 | Inositol transporter 4 | 0 |
| SG001_LG007CG010690 | 507 | inositol transporter 4-like | 0 | Inositol transporter 4 | 0 |
| ***RlIP5P7*** |  |  |  |  |  |
| SG001_LG004CG000150 | 592 | type IV inositol polyphosphate 5-phosphatase 7-like | 0 | Type IV inositol polyphosphate 5-phosphatase 7 | 0 |
| SG001_LG006CG031450 | 592 | type IV inositol polyphosphate 5-phosphatase 7-like | 0 | Type IV inositol polyphosphate 5-phosphatase 7 | 0 |
| ***RlLAP2*** |  |  |  |  |  |
| SG001_LG001CG019360 | 143 | leucine aminopeptidase 1-like | 1.71E−50 | Leucine aminopeptidase 1 | 8.39E−33 |
| SG001_LG002CG001050 | 576 | leucine aminopeptidase 1-like | 0 | Leucine aminopeptidase 2, chloroplastic | 0 |
| SG001_LG002CG001140 | 525 | leucine aminopeptidase 1-like | 0 | Leucine aminopeptidase 2, chloroplastic | 0 |
| SG001_LG002CG001150 | 576 | leucine aminopeptidase 1-like | 0 | Leucine aminopeptidase 2, chloroplastic | 0 |
| SG001_LG004CG004570 | 143 | leucine aminopeptidase 1-like | 1.66E−51 | Leucine aminopeptidase 1 | 3.50E−33 |
| SG001_LG007CG018780 | 143 | leucine aminopeptidase 1-like | 1.66E−51 | Leucine aminopeptidase 1 | 3.50E−33 |

**Table S8.** *Continued*.

| **Gene ID** | **Protein**  **length** | **NCBI nr blastp** |  | **UniProtKB/Swiss-Prot blastp** |  |
| --- | --- | --- | --- | --- | --- |
|  |  | **Gene description** | **E-value** | **Gene description** | **E-value** |
| ***RlLRK10L1.2*** |  |  |  |  |  |
| SG001_LG004CG035040 | 528 | leaf rust 10 disease-resistance locus receptor-like protein kinase-like 1.2 isoform X1 | 0 | leaf rust 10 disease-resistance locus receptor-like protein kinase-like 1.2 | 2.41E−90 |
| SG001_LG004CG035050 | 686 | leaf rust 10 disease-resistance locus receptor-like protein kinase-like 1.2 isoform X1 | 0 | leaf rust 10 disease-resistance locus receptor-like protein kinase-like 1.2 | 0 |
| SG001_LG007CG006950 | 661 | leaf rust 10 disease-resistance locus receptor-like protein kinase-like 1.2 isoform X2 | 0 | leaf rust 10 disease-resistance locus receptor-like protein kinase-like 1.2 | 0 |
| SG001_LG007CG006970 | 694 | leaf rust 10 disease-resistance locus receptor-like protein kinase-like 1.2 isoform X1 | 0 | leaf rust 10 disease-resistance locus receptor-like protein kinase-like 1.2 | 0 |
| SG001_LG007CG007030 | 662 | leaf rust 10 disease-resistance locus receptor-like protein kinase-like 1.2 isoform X2 | 0 | leaf rust 10 disease-resistance locus receptor-like protein kinase-like 1.2 | 0 |
| SG001_LG007CG007040 | 665 | leaf rust 10 disease-resistance locus receptor-like protein kinase-like 1.2 isoform X2 | 0 | leaf rust 10 disease-resistance locus receptor-like protein kinase-like 1.2 | 0 |
| SG001_LG007CG024200 | 681 | leaf rust 10 disease-resistance locus receptor-like protein kinase-like 1.2 isoform X2 | 0 | leaf rust 10 disease-resistance locus receptor-like protein kinase-like 1.2 | 0 |
| ***RlLRK91*** |  |  |  |  |  |
| SG001_LG002CG020490 | 559 | L-type lectin-domain containing receptor kinase IX.1-like | 0 | L-type lectin-domain containing receptor kinase IX.1 | 0 |
| SG001_LG002CG020520 | 650 | L-type lectin-domain containing receptor kinase IX.1-like | 0 | L-type lectin-domain containing receptor kinase IX.1 | 0 |
| SG001_LG002CG020530 | 687 | L-type lectin-domain containing receptor kinase IX.1-like | 0 | L-type lectin-domain containing receptor kinase IX.1 | 0 |
| SG001_LG002CG022320 | 648 | L-type lectin-domain containing receptor kinase IX.1-like | 0 | L-type lectin-domain containing receptor kinase IX.1 | 0 |
| SG001_LG002CG022330 | 636 | L-type lectin-domain containing receptor kinase IX.1-like | 0 | L-type lectin-domain containing receptor kinase IX.1 | 0 |
| SG001_LG003CG007100 | 664 | L-type lectin-domain containing receptor kinase IX.1-like | 0 | L-type lectin-domain containing receptor kinase IX.1 | 9.97E−180 |
| SG001_LG003CG007130 | 667 | L-type lectin-domain containing receptor kinase IX.1-like | 0 | L-type lectin-domain containing receptor kinase IX.1 | 7.88E−178 |
| SG001_LG003CG007140 | 700 | L-type lectin-domain containing receptor kinase IX.1-like | 0 | L-type lectin-domain containing receptor kinase IX.1 | 0 |
| SG001_LG003CG007150 | 528 | L-type lectin-domain containing receptor kinase IX.1-like | 0 | L-type lectin-domain containing receptor kinase IX.1 | 3.74E−132 |
| SG001_LG003CG007160 | 677 | L-type lectin-domain containing receptor kinase IX.1-like | 0 | L-type lectin-domain containing receptor kinase IX.1 | 1.97E−175 |
| SG001_LG004CG012890 | 702 | L-type lectin-domain containing receptor kinase IX.1-like | 0 | L-type lectin-domain containing receptor kinase IX.1 | 0 |
| SG001_LG004CG012920 | 538 | L-type lectin-domain containing receptor kinase IX.1-like | 0 | L-type lectin-domain containing receptor kinase IX.1 | 1.50E−157 |
| SG001_LG004CG012940 | 715 | L-type lectin-domain containing receptor kinase IX.1-like | 0 | L-type lectin-domain containing receptor kinase IX.1 | 0 |
| SG001_LG004CG013150 | 690 | L-type lectin-domain containing receptor kinase IX.1-like | 0 | L-type lectin-domain containing receptor kinase IX.1 | 0 |
| SG001_LG005CG023470 | 702 | L-type lectin-domain containing receptor kinase IX.1-like | 0 | L-type lectin-domain containing receptor kinase IX.1 | 2.67E−170 |
| SG001_LG005CG023690 | 679 | L-type lectin-domain containing receptor kinase IX.1-like | 0 | L-type lectin-domain containing receptor kinase IX.1 | 5.43E−131 |
| SG001_LG005CG023700 | 689 | L-type lectin-domain containing receptor kinase IX.1-like | 0 | L-type lectin-domain containing receptor kinase IX.1 | 0 |
| SG001_LG005CG023740 | 695 | L-type lectin-domain containing receptor kinase IX.1-like | 0 | L-type lectin-domain containing receptor kinase IX.1 | 0 |
| SG001_LG005CG023760 | 706 | L-type lectin-domain containing receptor kinase IX.1-like | 0 | L-type lectin-domain containing receptor kinase IX.1 | 0 |
| SG001_LG005CG023770 | 698 | L-type lectin-domain containing receptor kinase IX.1-like | 0 | L-type lectin-domain containing receptor kinase IX.1 | 0 |
| SG001_LG005CG023790 | 649 | L-type lectin-domain containing receptor kinase IX.1-like | 0 | L-type lectin-domain containing receptor kinase IX.1 | 2.10E−180 |
| SG001_LG005CG023810 | 681 | L-type lectin-domain containing receptor kinase IX.1-like | 0 | L-type lectin-domain containing receptor kinase IX.1 | 1.48E−169 |
| SG001_LG005CG027400 | 699 | L-type lectin-domain containing receptor kinase IX.1-like | 0 | L-type lectin-domain containing receptor kinase IX.1 | 0 |
| SG001_LG005CG030230 | 662 | L-type lectin-domain containing receptor kinase IX.1-like | 0 | L-type lectin-domain containing receptor kinase IX.1 | 1.29E−175 |
| SG001_LG005CG030250 | 661 | L-type lectin-domain containing receptor kinase IX.1-like | 0 | L-type lectin-domain containing receptor kinase IX.1 | 1.14E−130 |
| SG001_LG005CG030270 | 695 | L-type lectin-domain containing receptor kinase IX.1-like | 0 | L-type lectin-domain containing receptor kinase IX.1 | 0 |
| SG001_LG005CG030280 | 608 | L-type lectin-domain containing receptor kinase IX.1-like | 0 | L-type lectin-domain containing receptor kinase IX.1 | 9.74E−144 |
| SG001_Un031CG000010 | 630 | L-type lectin-domain containing receptor kinase IX.1-like | 0 | L-type lectin-domain containing receptor kinase IX.1 | 0 |

**Table S8.** *Continued*.

| **Gene ID** | **Protein**  **length** | **NCBI nr blastp** |  | **UniProtKB/Swiss-Prot blastp** |  |
| --- | --- | --- | --- | --- | --- |
|  |  | **Gene description** | **E-value** | **Gene description** | **E-value** |
| ***RlLRX4*** |  |  |  |  |  |
| SG001_LG002CG005780 | 662 | leucine-rich repeat extensin-like protein 4 | 0 | Leucine-rich repeat extensin-like protein 5 | 1.66E−156 |
| SG001_LG002CG005790 | 724 | leucine-rich repeat extensin-like protein 4 | 0 | Leucine-rich repeat extensin-like protein 5 | 0 |
| SG001_LG006CG040500 | 717 | leucine-rich repeat extensin-like protein 4 | 0 | Leucine-rich repeat extensin-like protein 5 | 0 |
| SG001_LG006CG051460 | 523 | leucine-rich repeat extensin-like protein 3 | 0 | Leucine-rich repeat extensin-like protein 4 | 2.21E−176 |
| ***RlPYL9*** |  |  |  |  |  |
| SG001_LG001CG029330 | 184 | abscisic acid receptor PYL9 | 9.40E−133 | Abscisic acid receptor PYL9 | 4.77E−106 |
| SG001_LG001CG029420 | 184 | abscisic acid receptor PYL9 | 9.40E−133 | Abscisic acid receptor PYL9 | 4.77E−106 |
| ***RlRbohA*** |  |  |  |  |  |
| SG001_LG007CG005060 | 790 | respiratory burst oxidase homolog protein A-like isoform X1 | 0 | Respiratory burst oxidase homolog protein F | 0 |
| SG001_LG007CG015390 | 953 | respiratory burst oxidase homolog protein A | 0 | Respiratory burst oxidase homolog protein F | 0 |
| ***RlRbohB*** |  |  |  |  |  |
| SG001_LG003CG006030 | 878 | respiratory burst oxidase homolog protein B | 0 | Respiratory burst oxidase homolog protein B | 0 |
| SG001_LG003CG006040 | 839 | respiratory burst oxidase homolog protein B | 0 | Respiratory burst oxidase homolog protein B | 0 |
| SG001_Un064CG000020 | 861 | respiratory burst oxidase homolog protein B | 0 | Respiratory burst oxidase homolog protein B | 0 |
| ***RlRbohC*** |  |  |  |  |  |
| SG001_LG007CG008750 | 926 | respiratory burst oxidase homolog protein C | 0 | Respiratory burst oxidase homolog protein C | 0 |
| SG001_LG007CG019090 | 764 | respiratory burst oxidase homolog protein C | 0 | Respiratory burst oxidase homolog protein C | 0 |
| ***RlRbohD*** |  |  |  |  |  |
| SG001_LG004CG029880 | 936 | respiratory burst oxidase homolog protein D | 0 | Respiratory burst oxidase homolog protein D | 0 |
| ***RlRbohE*** |  |  |  |  |  |
| SG001_LG002CG009230 | 923 | respiratory burst oxidase homolog protein E | 0 | Respiratory burst oxidase homolog protein E | 0 |
| SG001_LG002CG009450 | 907 | respiratory burst oxidase homolog protein E | 0 | Respiratory burst oxidase homolog protein E | 0 |
| SG001_LG002CG015870 | 662 | respiratory burst oxidase homolog protein E | 0 | Respiratory burst oxidase homolog protein E | 0 |
| ***RlRbohH*** |  |  |  |  |  |
| SG001_LG005CG045220 | 790 | putative respiratory burst oxidase homolog protein H isoform X1 | 0 | Putative respiratory burst oxidase homolog protein H | 0 |
| ***RlRSA1*** |  |  |  |  |  |
| SG001_LG002CG064630 | 93 | protein SHORT ROOT IN SALT MEDIUM 1 | 1.16E−47 | Protein SHORT ROOT IN SALT MEDIUM 1 | 7.36E−28 |
| SG001_LG004CG038240 | 1377 | protein SHORT ROOT IN SALT MEDIUM 1 | 0 | Protein SHORT ROOT IN SALT MEDIUM 1 | 3.87E−114 |
| SG001_LG005CG038770 | 107 | protein SHORT ROOT IN SALT MEDIUM 1 | 1.91E−58 | Protein SHORT ROOT IN SALT MEDIUM 1 | 2.99E−30 |
| SG001_LG006CG029800 | 93 | protein SHORT ROOT IN SALT MEDIUM 1 | 6.28E−48 | Protein SHORT ROOT IN SALT MEDIUM 1 | 2.46E−27 |
| ***RlSD25*** |  |  |  |  |  |
| SG001_LG002CG037570 | 815 | G-type lectin S-receptor-like serine/threonine-protein kinase SD2-5 | 0 | G-type lectin S-receptor-like serine/threonine-protein kinase SD2-5 | 0 |
| SG001_LG002CG037580 | 811 | G-type lectin S-receptor-like serine/threonine-protein kinase SD2-5 | 0 | G-type lectin S-receptor-like serine/threonine-protein kinase SD2-5 | 0 |
| SG001_LG002CG037670 | 815 | G-type lectin S-receptor-like serine/threonine-protein kinase SD2-5 | 0 | G-type lectin S-receptor-like serine/threonine-protein kinase SD2-5 | 0 |

**Table S8.** *Continued*.

| **Gene ID** | **Protein**  **length** | **NCBI nr blastp** |  | **UniProtKB/Swiss-Prot blastp** |  |
| --- | --- | --- | --- | --- | --- |
|  |  | **Gene description** | **E-value** | **Gene description** | **E-value** |
| ***RlSD25*** *(continued)* |  |  |  |  |  |
| SG001_LG002CG037680 | 821 | G-type lectin S-receptor-like serine/threonine-protein kinase SD2-5 | 0 | G-type lectin S-receptor-like serine/threonine-protein kinase SD2-5 | 0 |
| SG001_LG004CG000320 | 665 | G-type lectin S-receptor-like serine/threonine-protein kinase SD2-5 | 0 | G-type lectin S-receptor-like serine/threonine-protein kinase SD2-5 | 0 |
| SG001_LG006CG040040 | 812 | G-type lectin S-receptor-like serine/threonine-protein kinase SD2-5 | 0 | G-type lectin S-receptor-like serine/threonine-protein kinase SD2-5 | 0 |
| SG001_LG006CG040050 | 691 | G-type lectin S-receptor-like serine/threonine-protein kinase SD2-5 | 0 | G-type lectin S-receptor-like serine/threonine-protein kinase SD2-5 | 0 |
| ***RlSLAH3*** |  |  |  |  |  |
| SG001_LG001CG037640 | 610 | S-type anion channel SLAH2-like | 0 | S-type anion channel SLAH3 | 0 |
| SG001_LG001CG037650 | 606 | S-type anion channel SLAH2-like | 0 | S-type anion channel SLAH3 | 0 |
| SG001_LG001CG044970 | 571 | S-type anion channel SLAH2-like | 0 | S-type anion channel SLAH3 | 0 |
| SG001_LG001CG044980 | 610 | S-type anion channel SLAH2-like | 0 | S-type anion channel SLAH3 | 0 |
| SG001_LG001CG045000 | 590 | S-type anion channel SLAH2-like | 0 | S-type anion channel SLAH3 | 0 |
| SG001_LG001CG045010 | 610 | S-type anion channel SLAH2-like | 0 | S-type anion channel SLAH3 | 0 |
| SG001_LG001CG045520 | 564 | S-type anion channel SLAH2-like | 0 | S-type anion channel SLAH3 | 0 |
| SG001_LG001CG045570 | 610 | S-type anion channel SLAH2-like | 0 | S-type anion channel SLAH3 | 0 |
| SG001_LG001CG045580 | 590 | S-type anion channel SLAH2-like | 0 | S-type anion channel SLAH3 | 0 |
| SG001_LG001CG045590 | 610 | S-type anion channel SLAH2-like | 0 | S-type anion channel SLAH3 | 0 |
| ***RlSTRK1*** |  |  |  |  |  |
| SG001_LG006CG036850 | 331 | probable receptor-like protein kinase At1g80640 | 0 | Probable receptor-like protein kinase At1g80640 | 2.07E−25 |
| SG001_Un019CG000220 | 331 | probable receptor-like protein kinase At1g80640 | 0 | Probable receptor-like protein kinase At1g80640 | 2.07E−25 |
| ***RlSUV3*** |  |  |  |  |  |
| SG001_LG004CG000140 | 566 | DExH-box ATP-dependent RNA helicase DExH16, mitochondrial isoform X1 | 0 | DExH-box ATP-dependent RNA helicase DExH16, mitochondrial | 0 |
| SG001_LG006CG031460 | 572 | DExH-box ATP-dependent RNA helicase DExH16, mitochondrial isoform X1 | 0 | DExH-box ATP-dependent RNA helicase DExH16, mitochondrial | 0 |
| ***RlZEP*** |  |  |  |  |  |
| SG001_LG002CG017400 | 626 | zeaxanthin epoxidase, chloroplastic | 0 | Zeaxanthin epoxidase, chloroplastic | 0 |
| SG001_LG002CG017410 | 660 | zeaxanthin epoxidase, chloroplastic | 0 | Zeaxanthin epoxidase, chloroplastic | 0 |
| SG001_LG002CG017440 | 642 | zeaxanthin epoxdase, chloroplastic | 0 | Zeaxanthin epoxidase, chloroplastic | 0 |
